# HDX-MS optimized approach to characterize nanobodies as tools for biochemical and structural studies of class IB phosphoinositide 3-kinases

**DOI:** 10.1101/2021.06.01.446614

**Authors:** Manoj K Rathinaswamy, Kaelin D Fleming, Udit Dalwadi, Els Pardon, Noah J Harris, Calvin K Yip, Jan Steyaert, John E Burke

**Author notes:** To whom correspondence should be addressed: John E. Burke. These authors contributed equally Lead contact: John E Burke.

## Abstract

There is considerable interest in developing antibodies as modulators of signaling pathways. One of the most important signaling pathways in higher eukaryotes is the phosphoinositide 3-kinase (PI3K) pathway, which plays fundamental roles in growth, metabolism and immunity. The class IB PI3K, PI3K*γ*, is a heterodimeric complex composed of a catalytic p110*γ* subunit bound to a p101 or p84 regulatory subunit. PI3K*γ* is a critical component in multiple immune signaling processes and is dependent on activation by Ras and GPCRs to mediate its cellular roles. Here we describe the rapid and efficient characterization of multiple PI3K*γ* single chain camelid nanobodies using hydrogen deuterium exchange mass spectrometry (HDX-MS) for structural and biochemical studies. This allowed us to identify nanobodies that stimulated lipid kinase activity, blocked Ras activation and specifically inhibited p101-mediated GPCR activation. Overall, this reveals novel insight into PI3K*γ* regulation and identifies sites that may be exploited for therapeutic development.

**Highlights:**

– HDX-MS rapidly identifies epitopes of camelid single-chain nanobodies raised against Class IB PI3K complexes, p110*γ*/p101 and p110*γ*/p84
– A nanobody targeting p101 improves local resolution in EM studies with p110*γ*/p101 facilitating structural characterization of the complex
– Nanobodies that bind at the interfaces with the lipidated activators Ras and G*βγ* can prevent activation of p110*γ*/p101 and p110*γ*/p84

## Introduction

Class I Phosphoinositide-3-Kinases (PI3Ks) are important lipid signaling proteins which are frequently mis-regulated in human disease (Fruman et al., 2017). These enzymes produce phosphatidylinositol-3,4,5-trisphosphate at the plasma membrane, which recruits numerous downstream effectors that control growth, survival, metabolism and immunity (Madsen and Vanhaesebroeck, 2020). All class I PI3Ks are hetero-dimeric complexes composed of a p110 catalytic subunit bound to a regulatory subunit. The regulatory subunits are crucial in the activation of class I PI3Ks downstream of membrane localized signaling molecules including receptor tyrosine kinases (RTKs), G-protein coupled receptors (GPCRs) and the Ras family of small GTPases (Burke and Williams, 2015). The class IB PI3Ks are composed of a single catalytic subunit (p110*γ*) bound to one of two regulatory subunits, p101 or p84 (also called p87). Both PI3K*γ* complexes have important roles in immune cell migration, cytokine production and cardiac function (Hawkins and Stephens, 2015) and are therapeutic targets in cancer immunotherapy (De Henau et al., 2016; Kaneda et al., 2016) and inflammation (Camps et al., 2005). The regulatory subunits mediate activation of p110*γ*, with either p101 or p84 needed for activation by G*βγ*, derived from GPCRs (Maier et al., 1999; Stephens et al., 1997). The p101 subunit is unique as it has a direct binding site for G*βγ* (Vadas et al., 2013), while p84 merely potentiates G*βγ* binding to p110*γ*. Both complexes of p110*γ* are activated by Ras (Kurig et al., 2009), which is mediated by the Ras binding domain (RBD) of p110*γ* (Pacold et al., 2000).

These differences in regulation translate into distinct cellular functions with p110*γ*/p101 controlling immune cell migration while p110*γ*/p84 controls mast cell degranulation and production of reactive oxides from neutrophils (Bohnacker et al., 2009; Deladeriere et al., 2015). A full understanding of how PI3K*γ* is regulated has been hampered by lack of molecular details on complex assembly, and activation by GPCRs. In addition, due to the severe side effects of pan-PI3K ATP-competitive inhibitors, targeting specific PI3K*γ* complexes could be useful for therapeutic approaches in cancer and inflammatory diseases. The development of potent and specific biomolecules that modulate PI3K*γ* activity will be useful for structural, biochemical and cellular studies of PI3K regulation.

One of the most powerful biomolecules for optimizing structural studies and modulating signaling pathways are single chain antibodies from camelids, known as nanobodies (Hamers-Casterman et al., 1993). Nanobodies are the variable domains of the heavy-chain only camelid antibodies (V_HH_), and lack hydrophobic residues that would normally pack against the light chain (V_L_) in conventional antibodies. As a result, nanobodies can be expressed in high yield in multiple expression systems (Muyldermans, 2013). Nanobodies have an enlarged antigen binding surface with a longer CDR3 loop, which normally packs against the V_L_ in conventional dual chain antibodies (Desmyter et al., 1996). This coupled with their small size (∼15 kDa versus 150 kDa for conventional antibodies) provide nanobodies the potential to bind specifically to epitopes that are inaccessible for conventional antibodies, with high affinity. These advantages have resulted in their widespread use in research, testing and therapy (Uchański et al., 2020). Nanobodies have proven to be exceptional tools for optimizing structural biology approaches of protein assemblies. In X-ray crystallography, they can stabilize flexible protein regions, prevent aggregation/oligomerization, and offer novel crystal contact sites (Baranova et al., 2012; Domanska et al., 2011; Korotkov et al., 2009; Schubert et al., 2017). Nanobodies are powerful tools to lock protein complexes into specific, functionally-relevant conformational states. For example, nanobodies have provided insight into active/inactive states of receptors or different stages in the transport cycle of membrane channels and pumps (Huang et al., 2015; Rasmussen et al., 2011a; 2011b; Ruprecht et al., 2019; Smirnova et al., 2015). They have played an important role in understanding PI3K biology, as a specific nanobody was crucial in the crystallization of the 385 kDa class III PI3K complex (Rostislavleva et al., 2015). In addition to crystallography, they have been useful in electron microscopy to assist structural characterization and in the labelling of protein subunits in large complexes (García-Nafría et al., 2018; Laverty et al., 2019; Westfield et al., 2011).

In addition to their utilization in structural biology, nanobodies can be used to dissect and target signaling events in cells and organisms (Bannas et al., 2017; Beghein and Gettemans, 2017). They are a potent tool to interfere with protein-protein interactions *in vivo,* which has potential applications in multiple human diseases. This is highlighted by their utilization as neutralizing agents in betacoronavirus infection, including SARS CoV-1/2 and MERS CoV, through blocking the interaction of the viral spike protein with ACE2 (Huo et al., 2020; Wrapp et al., 2020). Nanobodies have been particularly powerful in modulating the signaling inputs and outputs of GPCRs (Manglik et al., 2017; Pardon et al., 2018). Conformationally selective nanobodies that bind to GPCRs can modulate agonist binding, receptor phosphorylation, and the recruitment of downstream partners including both G-proteins and *β*-arrestin (McMahon et al., 2020; Staus et al., 2016; Wingler et al., 2019). GPCR activation of G-protein signaling stimulates the dissociation of the heterotrimeric G protein into G*α*-GTP and a G*βγ* dimer, and nanobodies targeting the effector binding surface of G*βγ* inhibit G*βγ* signaling (Gulati et al., 2018). Development of nanobodies that disrupt specific inputs from GPCRs and other membrane receptors into distinct signaling pathways will be powerful in dissecting the molecular mechanisms of signaling and in developing novel therapeutics. Critical to determining the usefulness of nanobodies as structural chaperones and signaling modulators is the ability to rapidly determine their binding sites and how they might alter protein conformational dynamics.

Here we report the characterization of multiple PI3K*γ* binding nanobodies, and describe their application for both structural biology approaches and modulation/inhibition of the activation and kinase activity of PI3K*γ*. We utilized hydrogen deuterium exchange mass spectrometry (HDX-MS) to rapidly identify binding epitopes in p110*γ*, p101, and p84. This allowed us to identify and characterize p110*γ* binding nanobodies that activate lipid kinase activity, block activation by Ras, and a p101 binding nanobody that disrupts GPCR activation. Nanobodies were identified that stabilized flexible regions/domains, allowing for high resolution cryo electron microscopy (cryo-EM) studies (details described in a separate study). Overall, this work provides an HDX-MS enabled flow-path for the rapid characterization of nanobodies for structural and biochemical studies (Figure 1).

**Figure 1.**
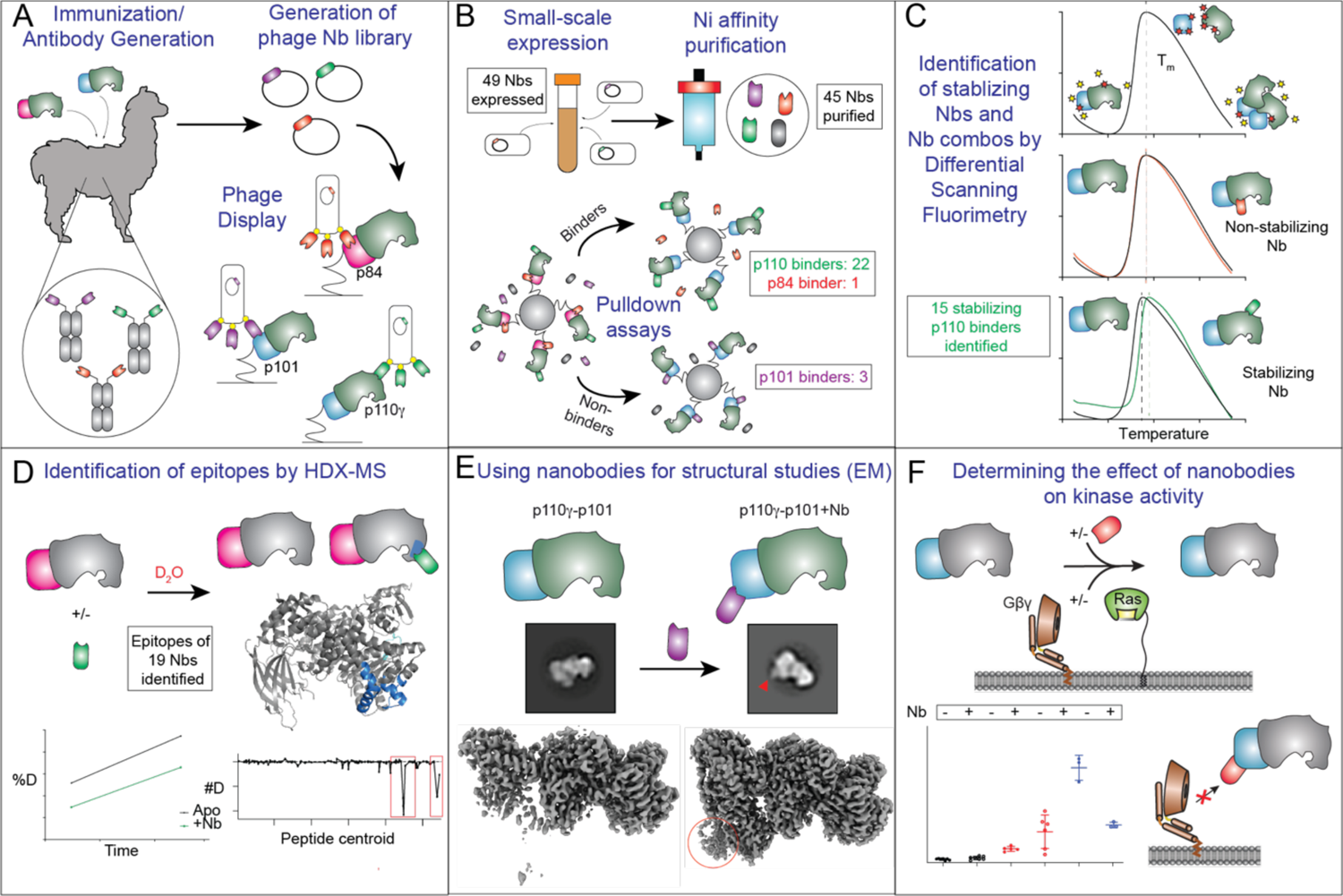
Schematic describing the flow-path for characterizing nanobodies that bind PI3Kγ as tools for structural and biochemical analysis. A. Isolation of nanobodies through immunization with PI3K*γ* complexes, p110*γ*-p101 and p110*γ*-p84 and nanobody selection by phage display. B. Small scale expression and purification of nanobodies for pulldown assays to select for binders to p110*γ*, p101 and p84 C. Differential scanning fluorimetry (DSF) to obtain nanobodies and nanobody combinations with stabilizing effects D. Identification of nanobody epitopes on p110*γ*, p101 and p84 by hydrogen-deuterium exchange mass spectrometry (HDX-MS) E. Utilizing nanobodies in electron microscopy to label PI3K*γ* subunits and to facilitate high resolution structural studies F. Utilizing nanobodies as tools to modulate PI3K*γ* regulation in kinase activity assays

## Results

### Characterization and identification of a panel of PI3Kγ nanobodies

Llamas were immunized with either the p110*γ*/p101 or p110*γ*/p84 complex and putative binders were identified using phage display from the B-cells as previously described (Pardon et al., 2014) (Fig. 1A). We identified 88 potential binders, which were classified according to the sequence of the third complementarity determining region (CDR3), into 49 families. Representative nanobodies from all families were recombinantly expressed in WK6 *E.coli* cells and purified using Nickel affinity chromatography. Members of four families could not be expressed, leading to 45 purified nanobodies. Streptavidin pulldown assays were performed using strep-tagged p110*γ*/p101 and p110*γ*/p84 complexes with the purified nanobodies to determine binding. To identify nanobodies that bound to p110*γ*, p101, or p84 we carried out pull downs on both p110*γ*/p101 and p110*γ*/p84. Nanobodies that bound both complexes were assumed to bind p110*γ*, while ones that bound specifically to either p110*γ*/p101 or p110*γ*/p84 were assumed to be p101 or p84 binders. Twenty-six nanobodies were identified as positive hits, of which twenty-two bound to p110*γ*, three bound to p101, and one to p84 (Fig. 1B, Table S1, and Source data). We used differential scanning fluorimetry (DSF) with nanobodies bound to PI3K*γ* to identify possible stabilizing effects. DSF measures the unfolding of proteins as a function of temperature, which allows for the identification of stabilizing binding partners by observing increases in the melting temperature (T_m_) (Fig. 1C). Fifteen nanobodies showed higher T_m_ values, including eight nanobodies which induced differences of 0.5°C or greater, indicating significant stabilizing effects (Table S1).

### HDX-MS enabled identification of nanobody binding epitopes

The p110*γ*/p101 and p110*γ*/p84 complexes were subjected to hydrogen deuterium exchange mass spectrometry (HDX-MS) in the presence of 19 different nanobodies confirmed by pulldowns, in order to determine the binding epitope, and possible differences in conformational dynamics. HDX-MS measures the exchange rate of hydrogens on the protein amide backbone with the deuterium from a deuterated buffer. These exchange rates are primarily dependent on protein secondary structure, and thus it is a powerful tool to examine protein conformational dynamics (Masson et al., 2017). The changes in HDX as a result, allow for the identification of potential binding interfaces or conformational changes induced upon complex formation with nanobodies. The full details of HDX data collection and analysis are shown in Table S2, with the full raw deuterium incorporation data, and differences in exchange with each nanobody included in the source data excel file.

HDX-MS was used to identify the epitopes of nineteen nanobodies, including the fifteen p110*γ* binding and four p101/p84 nanobodies (Fig. 1D). HDX experiments were carried out at two timepoints of H/D exchange (3 or 300 seconds at 18°C). The p110*γ*-binding nanobodies caused decreased exchange at regions spanning almost the entire p110*γ* sequence (Figure 2 and Figure S1+S2). The putative binding epitopes are indicated in the source data. Multiple nanobodies caused decreased exchange in the helical domain of p110*γ*, which has been observed for previous p110*γ*antibody and small molecule binding partners (Gangadhara et al., 2019; Rathinaswamy et al., 2021; Shymanets et al., 2015) due to the propagation of allosteric changes to the helical domain. Of the nanobodies studied by HDX, we were able to identity putative epitopes for the majority (Fig. 2, Fig S1+S2, Table S1). Of this group, we will focus the discussion on nanobodies that were further characterized, but the full raw data are included in the source data. The nanobody NB5-PIK3CG caused decreased HDX in a region spanning the end of the uncharacterized adaptor binding domain (ABD) and the first helix in the ABD-RBD linker (122-157)(Fig 2B). Nanobody NB7-PIK3CG caused decreased HDX throughout the helical domain, with the largest decrease being localized in the Ras binding domain (RBD) which is essential for Ras binding (196-211) (Fig 2C). The p101-binding nanobody, NB1-PIK3R5, induced large scale decreases in exchange within multiple regions in the final 200 amino acids, which has been identified as a critical region for G*βγ* activation (Vadas et al., 2013). The p101 region previously identified as the putative G*βγ* binding site (816-828) had a >25% protection in deuterium incorporation.

**Figure 2.**
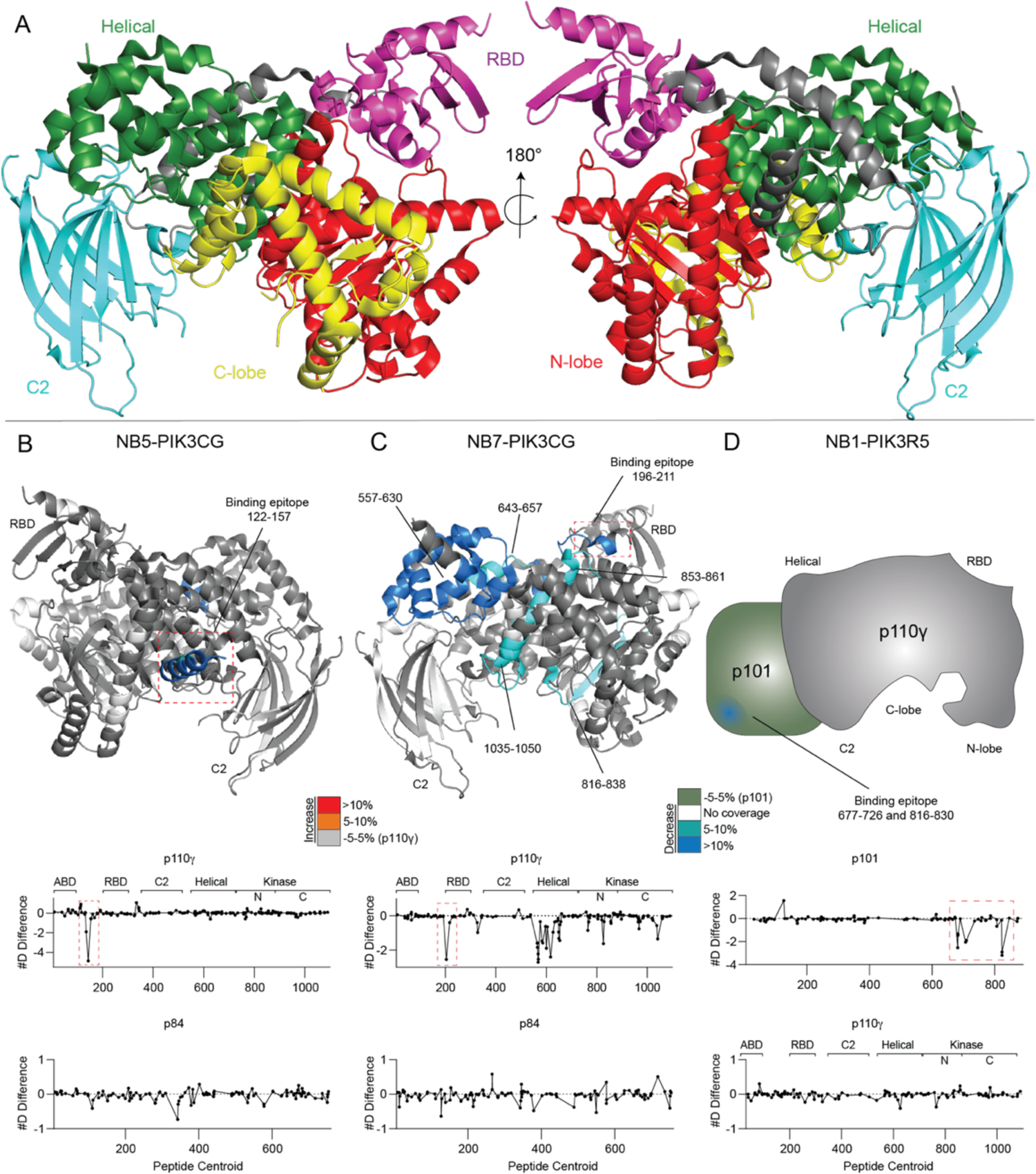
Using HDX-MS to identify nanobody epitopes on PI3Kγ. **A.** Domain organization of p110*γ*subunit with domains colored on the crystal structure of the 144-1102 crystal construct. (PDB ID: 6aud) **B.** HDX-MS differences in p110*γ*-p84 with the addition of NB5-PIK3CG mapped on a structural model of p110*γ*. The number of deuteron difference for all peptides analyzed over the entire deuterium exchange time course is shown for p110*γ* and p84. In panels **B-D**, peptides showing significant difference in deuterium exchange (>5%,>0.4 kDa) between conditions with and without nanobody are colored on the cartoon. **C.** HDX-MS differences in p110*γ*-p84 with the addition of NB7-PIK3CG mapped on a model of p110*γ*. The number of deuteron difference for all peptides analyzed over the entire deuterium exchange time course is shown for p110*γ* and p84. **D.** HDX-MS differences in p110*γ*-p101 with the addition of NB1-PIK3R5 mapped on a cartoon representation of p110*γ*/p101, as changes only occur in the p101 subunit, which has not been structurally characterized. The number of deuteron difference for all peptides analyzed over the entire deuterium exchange time course is shown for p110*γ* and p101.

### Nanobodies stabilize protein conformations enabling high resolution Cryo-EM analysis

Many nanobodies bound to regions of either p110*γ* or the p101 and p84 subunits that have not been characterized structurally up to this point. We first used utilized negative stain electron microscopy to analyze three nanobodies that bind novel regions of the p110*γ*-p101 complex: NB1-PIK3R5 (p101 C-terminus), NB2-PIK3R5 (p101) and NB5-PIK3CG (p110*γ* ABD) (Fig 3A). 2D analysis revealed that additional densities along the periphery of the complex corresponding to unique binding sites of these nanobodies. We were able to model the approximate location of these binding sites by integrating data from HDX-MS and negative stain EM (Fig. 3D). We also conducted cryo-EM analysis of the p110*γ*-p101 complex, generating a map at an overall resolution of 3.4 Å. However, this 3D reconstruction revealed that a large portion of p101 was poorly resolved in the EM density map, with this region matching the area where NB1-PIK3R5 bound based on the negative stain analysis and HDX-MS. Hence, we reconstituted a ternary complex of p110*γ*-p101 with NB1-PIK3R5 and vitrified this sample for cryo-EM analysis. We were able to obtain a 3D reconstruction at an overall resolution of 2.9 Å, with greatly improved local resolution compared to the apo complex at the NB1-PIK3R5 binding site (Fig. 3B+C) (the full details of this structural reconstruction are described in a separate manuscript). The improved quality of the EM density map resulting from the nanobody serves as a proof of principle for the HDX-MS guided approach to rapidly optimize nanobodies as structural chaperones. This enabled us to efficiently narrow down to the best possible nanobody to overcome the challenges facing structural investigations of p110*γ*-p101 (Fig 3D).

**Figure 3.**
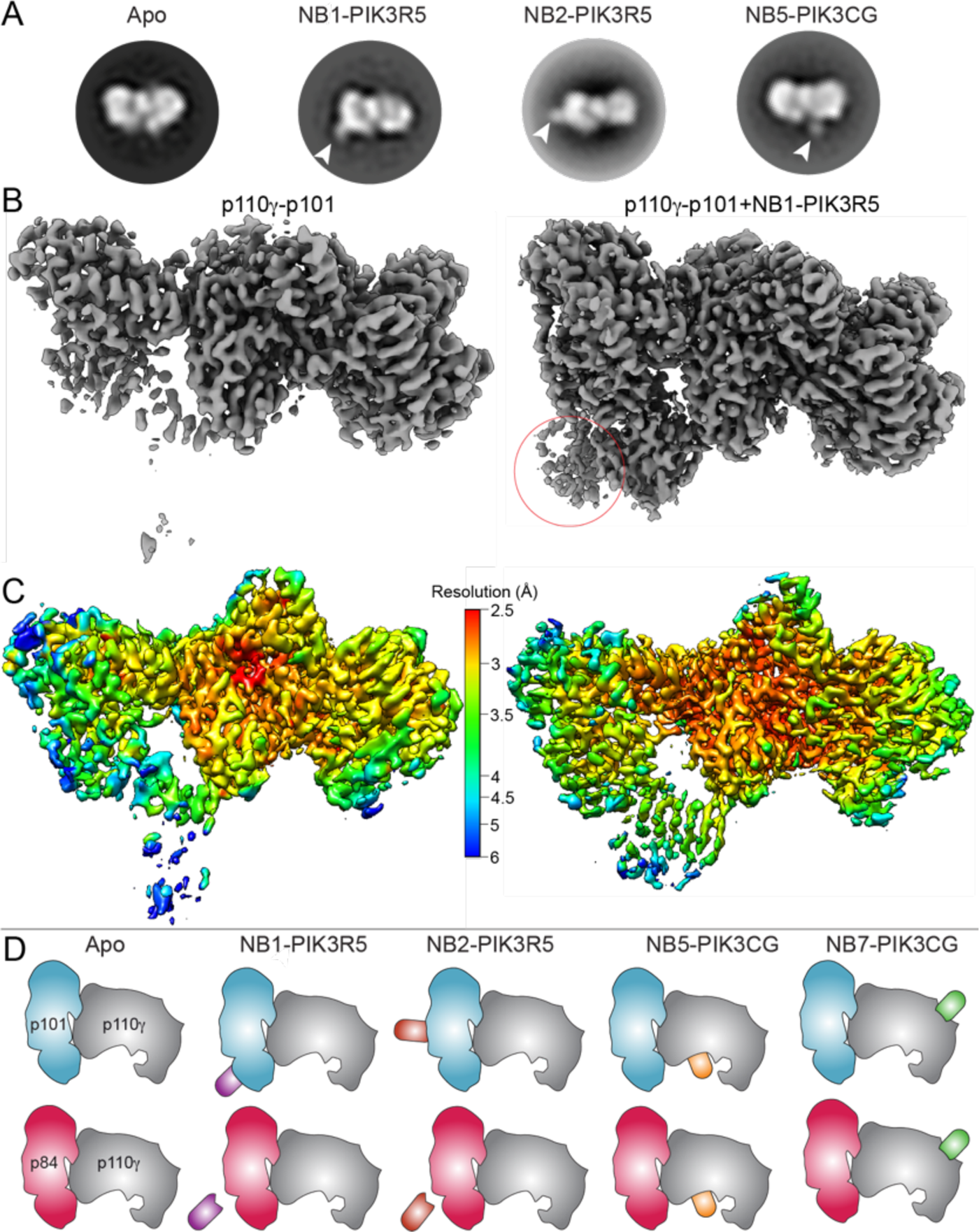
Nanobodies facilitate structural studies by EM. **A.** Representative 2D class averages of p110*γ*-p101 indicating positions of the nanobodies NB1-PIK3R5, NB2-PIK3R5 and NB5-PIK3CG as seen on negative stain EM **B.** Cryo-EM 3D reconstructions of p110*γ*-p101 and NB1-PIK3R5 bound p110*γ*-p101 showing the stabilizing effect of this nanobody. Density for the nanobody is circled. **C.** p110*γ*-p101 reconstructions with and without NB1-PIK3R5 colored according to local resolution as estimated using cryoSPARC v3.1 **D.** Cartoons showing approximate binding sites of NB1-PIK3R5, NB2-PIK3R5, NB5-PIK3CG and NB7-PIK3CG as determined by HDX-MS and negative stain EM.

*Nanobodies can both inhibit and activate* PI3K*γ lipid kinase activity*

The identification of nanobodies that bound at the protein interfaces for the upstream activators Ras (which binds the RBD of p110*γ*) and G*βγ* (binds the c-terminus of p101) led us to characterize their effects on lipid kinase activity and activation by lipidated Ras and G*βγ*. We hypothesized that nanobodies sharing the same binding epitopes as known PI3K*γ* activators could potentially disrupt PI3K*γ* activation and signaling. Additionally, the important role of the ABD domain in regulating class I PI3Ks, led to investigate the potential role of nanobodies binding this region. We selected three nanobodies for full biochemical characterization (the RBD binding nanobody NB5-PIK3CG, the p101 binding nanobody NB1-PIK3R5, and the ABD binding nanobody NB7-PIK3CG) using in vitro lipid kinase assays of its basal activity and activation by lipidated G*βγ* and Ras (Fig 1F, Fig 4). Assays were carried out with both the p110*γ*-p101 and p110*γ*-p84 complexes, to determine any complex specific modulatory effects. The development of biomolecules that can specifically target unique p110*γ* complexes would be an important tool to decipher PI3K*γ* signaling, as ATP competitive inhibitors will equally inhibit both complexes.

**Figure 4.**
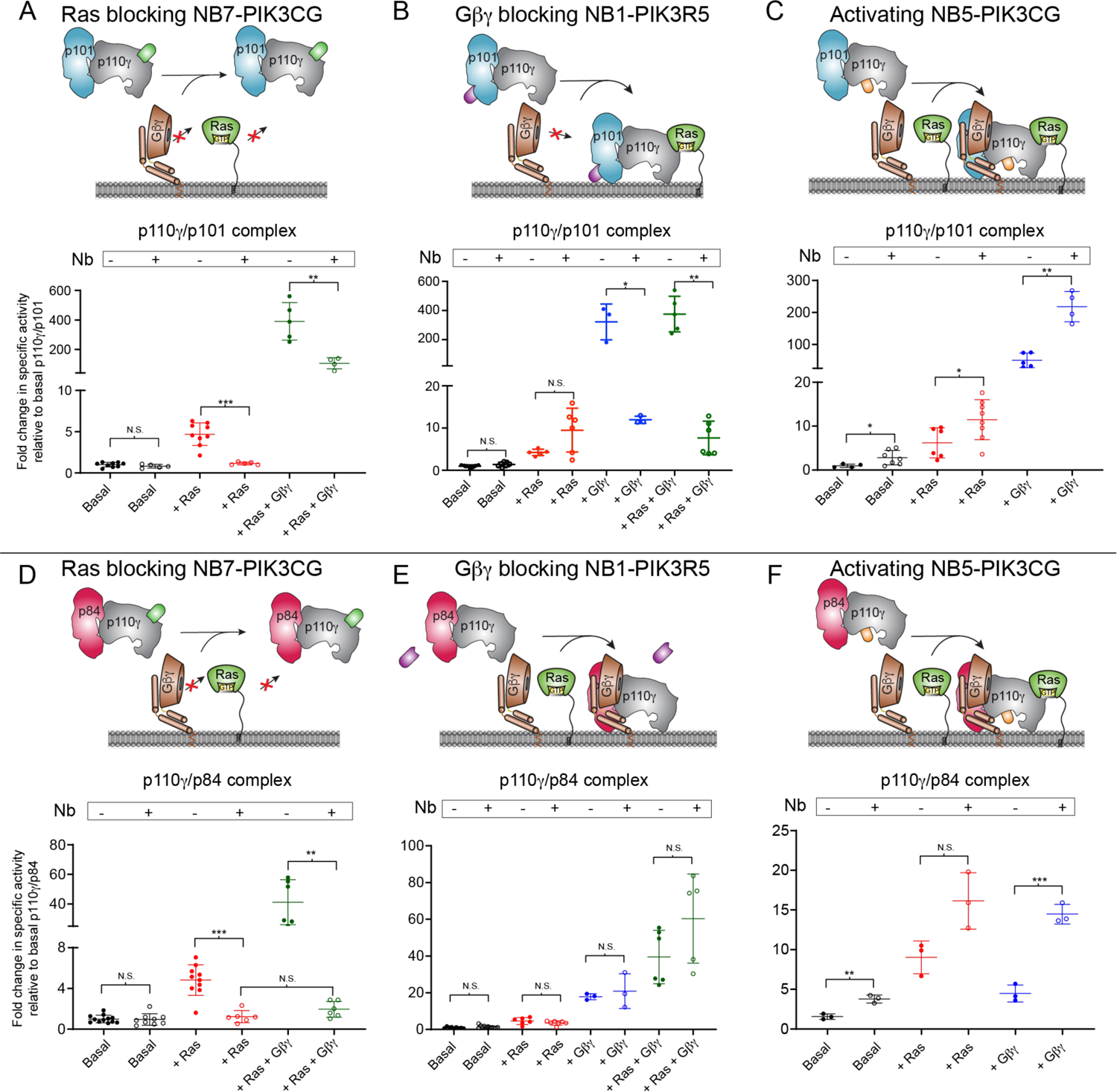
Nanobodies modulate PI3Kγ regulation. **A.** Lipid kinase activity assay showing the effect of NB7-PIK3CG binding on the specific activity of p110γ-p101 under the indicated conditions. For assays in **A-F**, plasma membrane mimic vesicles (20% phosphatidylserine (PS), 50% phosphatidylethanolamine (PE), 10% Cholesterol, 10% phosphatidylcholine (PC), 5% sphingomyelin (SM) and 5% phosphatidylinositol-3,4,5-trisphosphate (PIP2)) at 0.5 mg/mL final concentration were used. Final concentration of ATP was 100 μM. Final concentration of nanobody was 6 μM. Lipidated Gβγ and HRas G12V were present at 1.5 μM concentration. Two tailed p-values represented by the symbols as follows: ***<0.001; **<0.01; *<0.05; N.S.>0.05. **B.** Lipid kinase activity assay showing the effect of NB5-PIK3CG binding on the specific activity of p110*γ*-p101 under the indicated conditions. **C.** Lipid kinase activity assay showing the effect of NB1-PIK3R5 binding on the specific activity of p110*γ*-p101 under the indicated conditions. Biochemical assays in panels A-C were carried out with p110*γ*-p101 at 50-3,000 nM final concentration. **D.** Lipid kinase activity assay showing the effect of NB7-PIK3CG binding on the specific activity of p110*γ*-p84 under the indicated conditions. **E.** Lipid kinase activity assay showing the effect of NB5-PIK3CG binding on the specific activity of p110*γ*-p84 under the indicated conditions. **F.** Lipid kinase activity assay showing the effect of NB1-PIK3R5 binding on the specific activity of p110*γ*-p84 under the indicated conditions. Biochemical assays in panels D-F were carried out with p110*γ*-p84 at 1,500-3,000 nM final concentration.

The RBD-binding nanobody NB7-PIK3CG had no effect on basal lipid kinase activity of either p110*γ*-p101 or p110*γ*-p84 (Fig. 4A+D). For both complexes it completely blocked activation by lipidated Ras (Fig. 4A+D). This nanobody appeared to have a limited effect on G*βγ*activation of the p110*γ*-p101 complex, as it was still robustly activated, however, for the p110*γ*-p84 complex it caused complete disruption of both Ras and G*βγ* activation (Fig. 4A+D). This potentially could be utilized as a biased inhibitor that would preferentially inhibit p110*γ*-p84 over p110*γ*-p101. The contact site of the p101 binding nanobody NB1-PIK3R5 partially overlapped with the one identified for lipidated G*βγ*on membranes (Vadas et al., 2013). In light of this, we hypothesized that this nanobody would be able to specifically inhibit G*βγ*-mediated activation of p110*γ*/p101. G*βγ* activation of p110*γ*/p101 was almost completely inhibited in the presence of NB1-PIK3R5 (Fig 4B), with no effect on Ras activation. This nanobody caused no significant differences in lipid kinase activity under any conditions for the p110*γ*/p84 complex (Fig 4E). Due to its ability to potently inhibit only GPCR activation of the p110*γ*-p101 complex, NB1-PIK3R5 will be a powerful tool to selectively inhibit only p110*γ*-p101 over p110*γ*-p84 to decipher their specific roles in PI3K*γ* signaling.

We tested the effect of the p110*γ*ABD binding nanobody (NB5-PIK3CG) on lipid kinase activity. The ABD of p110*γ* is structurally uncharacterized, but the ABD domain of class IA PI3Ks is known to be a critical regulator of lipid kinase activity (Vadas et al., 2011). The NB5-PIK3CG nanobody activated lipid kinase activity under all conditions tested for both p110*γ*-p101 and p110*γ*-p84 (Fig. 3C+F). This reveals an unexpected and previously undescribed role of the ABD as a regulator of p110*γ* signaling, with molecules targeting this region able to modulate lipid kinase activity.

## Discussion

The class I PI3Ks are master regulators of growth, metabolism, and immunity (Fruman et al., 2017). Mutations leading to activation of the PI3K pathway are the most frequent alterations in human cancer (Lawrence et al., 2014). Small molecule inhibitors of class I PI3Ks are in clinical and pre-clinical development for cancer, immune disorders, developmental disorders, and inflammatory disorders. Partially limiting this approach is the severe side effects associated with pan-PI3K ATP-competitive inhibitors (McPhail and Burke, 2020). Further development of biomolecules that can selectively modulate distinct PI3K isoforms and complexes will be critical in fully understanding PI3K signaling, and may prove useful as therapeutics for multiple human diseases. Here we describe an HDX-MS optimized flow-path for the rapid identification of a panel of class IB binding nanobodies. These identified nanobodies enabled high resolution structural studies, and selectively modulated different p110*γ* regulatory complexes.

Nanobodies can be powerful tools in preventing protein-protein interactions, and they are rapidly entering the clinic for treatment of multiple diseases (Steeland et al., 2016), with the first nanobody drug Cablivi approved as a treatment for acquired thrombotic thrombocytopenic purpura (Muyldermans, 2021). In addition to this therapeutic role, one of the first applications of nanobodies was as chaperones facilitating structural biology approaches. This has enabled high resolution structures of multiple signaling proteins, highlighted by the foundational impact nanobodies have had on GPCR structural biology (Rasmussen et al., 2011a). A complication of generating optimized nanobodies, is the extensive screening required for biomolecules that enable either EM or X-ray approaches. Here we have described how HDX-MS can be utilized to rapidly identify epitopes for nanobodies and stabilizing conformational changes induced upon binding. HDX-MS is a well-established technique for efficiently defining antibody binding sites for biopharmaceuticals (Berkowitz et al., 2012), and has been used to define nanobody binding sites (Buckles et al., 2020; Rostislavleva et al., 2015). The identification of the NB1-PIK3R5 nanobody, which stabilized the dynamic C-terminus of the p101 subunit, allowed us to obtain a high resolution map of the p110*γ*-p101 complex by cryo-EM, which is described in depth in another manuscript. Overall, our HDX-MS based approach allows for a repeatable method to rapidly identify the most suitable nanobodies to optimize X-ray crystallography and cryo-EM approaches.

Our combined HDX-MS and EM structural studies revealed multiple nanobodies that bound at critical regulatory interfaces involved in the binding of Ras and G*βγ* in both p110*γ* and p101. We identified two nanobodies (NB6-PIK3CG + NB7-PIK3CG) that bound to the RBD in p110*γ*, which contains the Ras binding interface (Pacold et al., 2000). Membrane reconstitution assays of Ras activation showed that NB7-PIK3CG potently inhibited Ras activation for both the p110*γ*-p101 and p110*γ*-p84 complexes. Intriguingly, for p110*γ*-p84, this nanobody also disrupted activation by G*βγ*. The p110*γ*-p84 complex is proposed to strictly require Ras for activation *in vivo* (Kurig et al., 2009), however, this data suggests a key role of the RBD in the activation by both Ras and G*βγ* in this complex. NB7-PIK3CG induced large-scale allosteric changes within the helical domain of p110*γ* (Fig. 2C). It has been shown that the G*βγ* binding interface in p110*γ* lies within the helical domain (Vadas et al., 2013), and thus a potential molecular mechanism for this inhibition is the disruption of the helical domain-G*βγ* interaction through nanobody-induced allosteric changes. The p110*γ*-p101 complex has a second G*βγ* binding site in p101 (Rynkiewicz et al., 2020; Stephens et al., 1997) which potentially allows it to overcome this inhibitory effect and still be activated by G*βγ*. This difference between the two PI3K*γ* complexes will likely enable this nanobody to be used as a biased p110*γ*-p84 inhibitor. A similar effect was seen with a C2 binding antibody that was able to selectively inhibit GPCR activation of p110*γ*-p84 over p110*γ*-p101 (Shymanets et al., 2015). Together these biomolecules will be useful as tools to study the specific roles of p110*γ*-p84 in cell signaling.

The p110*γ*-p101 complex contains G*βγ* binding sites in both p110*γ* and p101 (Rynkiewicz et al., 2020; Shymanets et al., 2013; Stephens et al., 1997; Vadas et al., 2013), with the unique p101 G*βγ* binding site being an attractive target for the development of molecules to selectively inhibit GPCR activation of the p110*γ*-p101 complex. Nanobodies have been developed that can inhibit/modulate GPCR signaling at multiple levels (De Groof et al., 2019), including ones that target the extracellular (Scholler et al., 2017) and intracellular (Irannejad et al., 2013) (Rasmussen et al., 2011a) faces of GPCRs, as well as the released G*βγ* heterodimer (Gulati et al., 2018). The development of molecules that can specifically block activation of a single G*βγ*effector will have major advantages both as tools, and as potential therapeutics. The NB1-PIK3R5 nanobody bound to p101 at a site previously proposed as the G*βγ* binding site (Vadas et al., 2013), and selectively inhibited G*βγ*activation of only the p110*γ*-p101 complex. Selectively inhibiting the p110*γ*-p101 complex has potential applications and advantages in targeting p110*γ* in disease. In heart failure, the p110*γ*-p101 complex is upregulated, while the p110*γ*-p84 complex plays an important role in maintaining cardiac contractility (Patrucco et al., 2004; Perino et al., 2011), highlighting the potential advantage of specifically targeting p110*γ*-p101. Initiation of toll-like receptor (TLR) signaling activates p110*γ* through engagement with Rab8 (Luo et al., 2014), with this being proposed to selectively activate p110*γ*-p101 through an unknown mechanism (Luo et al., 2018). This might indicate an advantage of targeting p110*γ*-p101 in TLR driven inflammation. Further structural and biochemical optimization of biomolecules binding at this site in p101 may play an important role in understanding p110*γ* signaling and designing potential therapeutics.

Overall, this study using HDX-MS to probe a family of PI3K*γ* binding nanobodies identified a wide variety of biomolecules that were useful in both high-resolution structural analysis and as selective modulators of PI3K activity. This approach can be employed for other large multi-component protein complexes, and is uniquely well-suited to develop and identify biomolecules that can allosterically modulate enzyme activity outside of the active site.

## Acknowledgements

J.E.B. is supported the Canadian Institute of Health Research (CIHR, 168998), a Michael Smith Foundation for Health Research (MSFHR, scholar 17686), and the Cancer Research Society (CRS-24368). C.K.Y. is supported by CIHR (FDN-143228) and the Natural Sciences and Engineering Research Council of Canada (RGPIN-2018-03951). JS and EP acknowledge the support and the use of resources of Instruct-ERIC, part of the European Strategy Forum on Research Infrastructures (ESFRI), and the Research Foundation - Flanders (FWO) and thank Nele Buys for the technical assistance during Nanobody discovery. This research project was supported in part by the UBC High Resolution Macromolecular Cryo-Electron Microscopy Facility (HRMEM). A portion of this research was supported by NIH grant U24GM129547 and performed at the PNCC at OHSU and accessed through EMSL (grid.436923.9), a DOE Office of Science User Facility sponsored by the Office of Biological and Environmental Research. We appreciate help from Theo Humphreys and Rose Marie Haynes with data collection at PNCC.

## STAR METHODS

### Key Resource Table

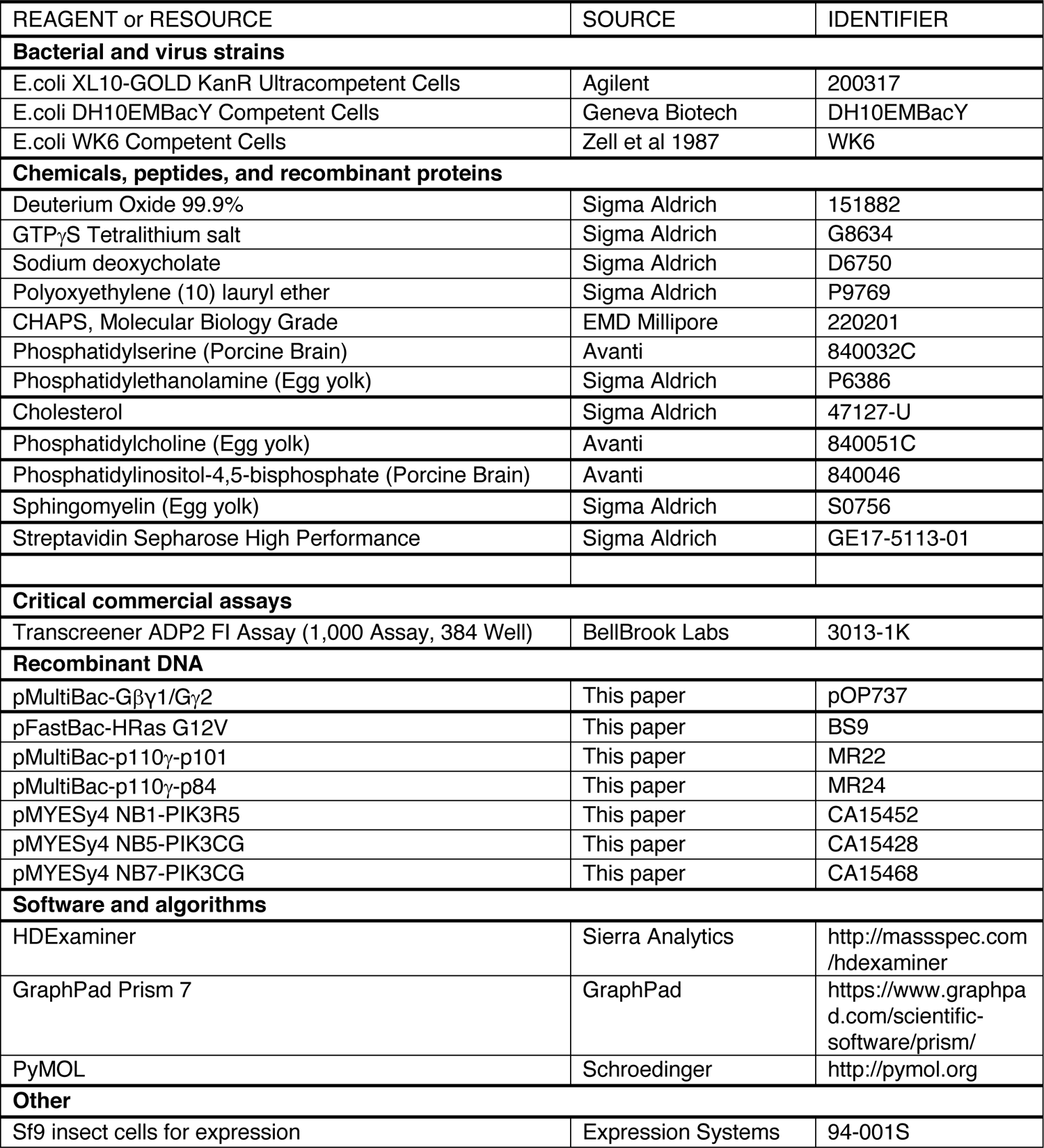

#### Virus Generation and Amplification

PI3Kγ complexes were expressed from a MutliBac plasmid with a 10X histidine tag, a 2X-strep tag and a Tobacco Etch Virus protease cleavage site on the N-terminus of the regulatory subunits. The plasmids encoding genes for insect cell expression were transformed into DH10MultiBac cells (MultiBac, Geneva Biotech) containing the baculovirus viral genome (bacmid) and a helper plasmid expressing transposase to transpose the expression cassette harbouring the gene of interest into the baculovirus genome. Bacmids with successful incorporation of the expression cassette into the bacmid were identified by blue-white screening and were purified from a single white colony using a standard isopropanol-ethanol extraction method. Briefly, colonies were grown overnight (16 hours) in 3-5 mL 2xYT (BioBasic #SD7019). Cells were pelleted by centrifugation and the pellet was resuspended in 300 μL P1 Buffer (50 mM Tris-HCl, pH 8.0, 10 mM EDTA, 100 mg/mL RNase A), chemically lysed by the addition of 300 μL Buffer P2 (1% sodium dodecyl sulfate (SDS) (W/V), 200 mM NaOH), and the lysis reaction was neutralized by addition of 400 μL Buffer N3 (3.0 M potassium acetate, pH 5.5). Following centrifugation at 21130 RCF and 4 °C (Rotor #5424 R), the supernatant was separated and mixed with 800 mL isopropanol to precipitate the DNA out of solution. Further centrifugation at the same temperature and speed pelleted the Bacmid DNA, which was then washed with 500 μL 70% Ethanol three times. The Bacmid DNA pellet was then dried for 1 minute and re-suspended in 50 mL Buffer EB (10 mM Tris-Cl, pH 8.5; All buffers from QIAprep Spin Miniprep Kit, Qiagen #27104). Purified bacmid was then transfected into Sf9 cells. 2 mL of Sf9 cells between 0.3-0.5X10^6^ cells/mL were aliquoted into the wells of a 6-well plate and allowed to attach, creating a monolayer of cells at ∼70-80% confluency. Transfection reactions were prepared by the addition of 2-10 ug of bacmid DNA to 100 μL 1xPBS and 12 mL polyethyleneimine (PEI) at 1 mg/mL (Polyethyleneimine ‘‘Max’’ MW 40.000, Polysciences #24765, USA) to 100 μL 1xPBS. The bacmid-PBS and the PEI-PBS solutions were mixed together, and the reaction occurred for 20-30 minutes before addition drop-by-drop to an Sf9 monolayer containing well. Transfections were allowed to proceed for 5-7 days before harvesting virus containing supernatant as a P1 viral stock.

#### Purification of PI3Kγ complexes

PI3Kγ catalytic and regulatory subunits were co-expressed in Spodoptera frugiperda (Sf9) cells using the baculovirus expression system. After 55 hours, the cells were harvested and the pellets were resuspended in lysis buffer (20 mM Tris pH 8.0, 100 mM NaCl, 10 mM imidazole pH 8.0, 2 mM beta-mercaptoethanol (βME), 5% (v/v) glycerol, Protease Inhibitor Cocktail (Millipore Protease Inhibitor Cocktail Set III, Animal-Free)) on ice. The resuspended pellets were sonicated for 2.5 minutes at level 4.0 with cycles consisting of 15 seconds ON/OFF using the Misonix Sonicator 3000. Triton X-100 was added to the cell lysate at a final concentration of 0.1% (v/v), and the lysates were centrifuged at 14,000 rpm at 4 °C for 45 minutes in a JA-20 rotor. The supernatant was loaded onto a HisTrap™ FF column (GE Healthcare Life Sciences) equilibrated with NiNTA A buffer (20 mM Tris pH 8.0, 100 mM NaCl, 10 mM imidazole pH 8.0, 5% (v/v) glycerol) and a high salt wash was performed using NiNTA A High Salt Buffer (20 mM Tris pH 8.0, 1 M NaCl, 10 mM imidazole pH 8.0, 5% (v/v) glycerol, 2mM βME). The protein was washed on an AKTA Start FPLC (GE) system with 4 column volumes (CV) of NiNTA A, 4 CV of 94% NiNTA A/6% NiNTA B (20 mM Tris pH 8.0, 100 mM NaCl, 200 mM imidazole pH 8.0, 2 mM βME, 5% (v/v) glycerol), and eluted in 2 CV of NiNTA B buffer. The eluted protein was loaded onto a StrepTrap™ column (GE) equilibrated with gel filtration buffer (GFB) (20 mM Tris pH 8.5, 100 mM NaCl, 50 mM (NH4)2SO4, 0.5 mM Tris (2-carboxyethyl) phosphine (TCEP)). For pulldowns, the column was washed with 2 CV GFB and the tagged protein was eluted in 2 CV GFB containing 2.5 mM Desthiobiotin. For studies using untagged protein, the washed column was incubated on ice overnight with 100 uL Tobacco Etch Virus Protease (1 mg/mL) diluted in GFB and eluted the following day. The protein was concentrated in an Amicon 50K Concentrator (Millipore Sigma). Gel filtration chromatography was performed on an AKTA Pure (GE) using a Superdex™ 200 10/300 Increase column (GE) equilibrated in GFB. The fractions containing the protein were pooled and concentrated, flash frozen in liquid nitrogen, and stored at −80 °C. All purification steps were analyzed using SDS-PAGE.

#### Nanobody generation and small scale nanobody purification

Nanobody discovery was carried out as previously described by the Steyaert lab at the VIB-VUB Center for Structural Biology (Pardon et al., 2014). Briefly, one llama was immunized 6 times with in total 900µg of p110*γ*/p101, another llama was immunized 6 times with in total 830µg of p110*γ*/p84 complex. Four days after the final boosts, peripheral blood lymphocytes were collected from both animals and RNA was extracted and used for creating cDNA encoding the ORFs of nanobodies. After PCR amplification, each library of Nanobody ORFs was cloned in the pMESy4 phage display vector (GenBank KF415192) creating 2 phage display libraries. Nanobodies were obtained after phage display selection on either p110*γ*/p101 or p110*γ*/p84 complex. In total 88 nanobodies were identified and classified into 49 families according to CDR3 sequence similarity. Nanobodies were expressed from pMYESy4 vectors in the periplasms of Escherichia coli WK6 cells. For pulldowns and HDX-MS, 5 mL cultures were grown to OD600 of 0.7 in Terrific Broth containing 0.1% glucose and 2mM MgCl2 in the presence of 100 μg/mL ampicillin and was induced with 0.5 mM isopropyl-β-D-thiogalactoside (IPTG). For assays and EM studies, nanobodies were expressed in 1L cultures. Cells were harvested the following day by centrifuging at 2500 RCF (Eppendorf Centrifuge 5810 R) and the pellet was snap-frozen in liquid nitrogen. Nanobodies were isolated from these pellets by osmotic shock. Cell pellets were resuspended TES buffer (3 mL for pellets from 5mL cultures or 15 mL for pellets from 1L cultures) (0.2 M Tris pH 8.0, 0.5 mM ethylenediaminetetraacetic acid (EDTA), 0.5 M sucrose) with Protease Inhibitor Cocktail Set III and rotated for 45 minutes at 4 °C. Twice the volume of TES/4 buffer was added to the cells and rotated for 30 minutes at 4 °C, followed by centrifugation at 14,000 rpm for 30 minutes at 4 °C in a JA-20 rotor. The periplasmic extract was loaded onto a HisTrap™ HP column (GE) equilibrated with NiNTA buffer (1X PBS pH 7.4, 1 M NaCl), followed by a wash with 5 CV NiNTA A wash buffer (1X PBS pH 7.4, 15 mM imidazole, 1 M NaCl). The protein was eluted with 2 CV of NiNTA B buffer (1X PBS pH 7.4, 200 mM imidazole, 1 M NaCl). The protein elution was exchanged into PI3K GFB with a 5 mL HiTrap™ Desalting column (GE). The success of purification was determined by SDS-PAGE.

#### Purification of lipidated Gβγ

Full length, lipidated human Gβγ (Gβ_1_γ_2_) was expressed in Sf9 insect cells and purified as described previously (Kozasa and Gilman, 1995). After 65 hours of expression, cells were harvested and the pellets were frozen as described above. Pellets were resuspended in lysis buffer (20 mM HEPES pH 7.7, 100 mM NaCl, 10 mM βME, protease inhibitor (Protease Inhibitor Cocktail Set III, Sigma)) and sonicated for 2 minutes (15s on, 15s off, level 4.0, Misonix sonicator 3000). The lysate was spun at 500 RCF (Eppendorf Centrifuge 5810 R) to remove intact cells and the supernatant was centrifuged again at 25,000 g for 1 hour (Beckman Coulter JA-20 rotor). The pellet was resuspended in lysis buffer and sodium cholate was added to a final concentration of 1% and stirred at 4°C for 1 hour. The membrane extract was clarified by spinning at 10,000 g for 30 minutes (Beckman Coulter JA-20 rotor). The supernatant was diluted 3 times with NiNTA A buffer (20 mM HEPES pH 7.7, 100 mM NaCl, 10 mM Imidazole, 0.1% C_12_E_10_, 10mM βME) and loaded onto a 5 mL HisTrap™ FF crude column (GE Healthcare) equilibrated in the same buffer. The column was washed with NiNTA A, 6% NiNTA B buffer (20 mM HEPES pH 7.7, 25 mM NaCl, 250 mM imidazole pH 8.0, 0.1% C12E10, 10 mM βME) and the protein was eluted with 100% NiNTA B. The eluent was loaded onto HiTrap^TM^ Q HP anion exchange column equilibrated in Hep A buffer (20 mM Tris pH 8.0, 8 mM CHAPS, 2 mM Dithiothreitol (DTT)). A gradient was started with Hep B buffer (20 mM Tris pH 8.0, 500 mM NaCl, 8 mM CHAPS, 2 mM DTT) and the protein was eluted in ∼50% Hep B buffer. The eluent was concentrated in a 30,000 MWCO Amicon Concentrator (Millipore) to < 1 mL and injected onto a Superdex^TM^ 75 10/300 GL size exclusion column (GE Healthcare) equilibrated in Gel Filtration buffer (20 mM HEPES pH 7.7, 100 mM NaCl, 10 mM CHAPS, 2 mM TCEP). Fractions containing protein were pooled, concentrated, aliquoted, frozen and stored at −80°C.

#### Expression Purification of Lipidated HRas G12V

Full-length HRas G12V was expressed by infecting 500 mL of Sf9 cells with 5 mL of baculovirus. Cells were harvested after 55 hours of infection and frozen as described above. The frozen cell pellet was resuspended in lysis buffer (50 mM HEPES pH 7.5, 100 mM NaCl, 10 mM βME and protease inhibitor (Protease Inhibitor Cocktail Set III, Sigma)) and sonicated on ice for 1 minute 30 seconds (15s ON, 15s OFF, power level 4.0) on the Misonix sonicator 3000. Triton-X 114 was added to the lysate to a final concentration of 1%, mixed for 10 minutes at 4°C and centrifuged at 25,000 rpm for 45 minutes (Beckman Ti-45 rotor). The supernatant was warmed to 37°C for few minutes until it turned cloudy following which it was centrifuged at 11,000 rpm at room temperature for 10 minutes (Beckman JA-20 rotor) to separate the soluble and detergent-enriched phases. The soluble phase was removed, and Triton-X 114 was added to the detergent-enriched phase to a final concentration of 1%. This phase separation was performed for a total of 3 times. Imidazole pH 8.0 was added to the detergent phase to a final concentration of 15 mM and the mixture was incubated with Ni-NTA agarose beads (Qiagen) for 1 hour at 4°C. The beads were washed with 5 column volumes of Ras-NiNTA buffer A (20mM Tris pH 8.0, 100mM NaCl, 15mM imidazole pH 8.0, 10mM βME and 0.5% Sodium Cholate) and the protein was eluted with 2 column volumes of Ras-NiNTA buffer B (20mM Tris pH 8.0, 100mM NaCl, 250mM imidazole pH 8.0, 10mM βME and 0.5% Sodium Cholate). The protein was buffer exchanged to Ras-NiNTA buffer A using a 10,000 kDa MWCO Amicon concentrator, where protein was concentrated to ∼1mL and topped up to 15 mL with Ras-NiNTA buffer A and this was repeated a total of 3 times. GTPγS was added in 2-fold molar excess relative to HRas along with 25 mM EDTA. After incubating for an hour at room temperature, the protein was buffer exchanged with phosphatase buffer (32 mM Tris pH 8.0, 200 mM Ammonium Sulphate, 0.1 mM ZnCl_2_, 10 mM βME and 0.5% Sodium Cholate). 1 unit of immobilized calf alkaline phosphatase (Sigma) was added per milligram of HRas along with 2-fold excess nucleotide and the mixture was incubated for 1 hour on ice. MgCl_2_ was added to a final concentration of 30 mM to lock the bound nucleotide. The immobilized phosphatase was removed using a 0.22-micron spin filter (EMD Millipore). The protein was concentrated to less than 1 mL and was injected onto a Superdex^TM^ 75 10/300 GL size exclusion column (GE Healthcare) equilibrated in gel filtration buffer (20 mM HEPES pH 7.7, 100 mM NaCl, 10 mM CHAPS, 1 mM MgCl2 and 2 mM TCEP). The protein was concentrated to 1 mg/mL using a 10,000 kDa MWCO Amicon concentrator, aliquoted, snap-frozen in liquid nitrogen and stored at −80°C.

#### Streptavidin Pulldown assays

To confirm binding, purified nanobodies at 3μM final concentration were mixed with tagged p110γ-p84 at 2μM final concentration in PI3K GFB and incubated for 15 minutes on ice. To this mixture, streptavidin beads (GE Healthcare) equilibrated in GFB was added and incubated again for 15 minutes. The beads were spun down at 500 g (Eppendorf centrifuge 5424 R) and the supernatant was removed. The beads were then washed for a total of three times in GFB. Following the final wash, the beads were spun down, the supernatant was removed and the proteins were eluted in GFB containing 2.5 mM desthiobiotin. The beads were spun down again following which the supernatant was mixed with loading dye and run on a 4-12% Nu-PAGE gel (Invitrogen: NP0321BOX). Binders to both p110γ and p84 were identified by the presence of bands corresponding to the nanobodies (∼15 kDa). Pull down assays were performed again on all nanobodies with tagged p110γ-p101 to identify binders to p101.

#### Differential Scanning Fluorimetry

Differential scanning fluorimetry was performed using the Applied Biosystems StepOnePlus™ RT-PCR instrument (ThermoFisher Scientific, cat. 4376600) with the excitation and emission wavelengths set to 587 and 607 nm, respectively. Briefly, p110γ-p101/ p110γ-p84 at a concentration of ∼1 μM was dispensed into a 96-well plate and mixed with ∼2 μM nanobody in PI3K GFB. SYPRO orange (Invitrogen) was diluted to 2.5x concentration, from a 5,000x stock. For thermal stability measurements, the temperature scan rate was fixed at 0.5 °C/min and the temperature range spanned 20 °C to 95 °C. Data analysis was performed using Protein Thermal Shift Software v1.4 (ThermoFisher Scientific, cat. 4466038), which determined melting temperatures (Tms) of individual replicates by fitting fluorescence data to a two-state Boltzman model.

### Hydrogen-Deuterium Exchange Mass Spectrometry (HDX-MS)

#### Preliminary HDX with p110γ-p101, and p110γ-p84 with nanobodies

HDX was performed by pre-incubating either p110γ-p101 or p110γ-p84 and 2-fold excess of nanobody for 2 minutes. After equilibration H/D exchange was carried out by dilution into a D2O buffer for either 3 or 300 seconds at 18 °C in a 50 ul reaction volume. The final total amount of PI3Kγ (with either p101 or p84) and nanobody were 20 pmol and 30 pmol, respectively. D2O buffer (20 mM HEPES pH 7.5, 100 mM NaCl, 96% D2O) was added to the protein samples to initiate hydrogen-deuterium exchange (final 84.5% D2O) and the reaction was quenched with an acidic quench solution (0.6 M guanidine-HCl, 0.9% formic acid final). The samples were immediately frozen in liquid nitrogen at −80°C.

#### Protein digestion and MS/MS data collection

Protein samples were rapidly thawed and injected onto an integrated fluidics system containing a HDx-3 PAL liquid handling robot and climate-controlled chromatography system (LEAP Technologies), a Dionex Ultimate 3000 UHPLC system, as well as an Impact HD QTOF Mass spectrometer (Bruker). The protein was run over either one (at 10°C) or two (at 10°C and 2°C) immobilized pepsin columns (Applied Biosystems; Poroszyme Immobilized Pepsin Cartridge, 2.1 mm x 30 mm; Thermo-Fisher 2-3131-00; Trajan; ProDx protease column, 2.1 mm x 30 mm PDX.PP01-F32) at 200 mL/min for 3 minutes. The resulting peptides were collected and desalted on a C18 trap column (Acquity UPLC BEH C18 1.7mm column (2.1 x 5 mm); Waters 186003975). The trap was subsequently eluted in line with a C18 reverse-phase separation column (Acquity 1.7 mm particle, 100 x 1 mm^2^ C18 UPLC column, Waters 186002352), using a gradient of 3-35% B (Buffer A 0.1% formic acid; Buffer B 100% acetonitrile) over 11 minutes immediately followed by a gradient of 35-80% over 5 minutes. Mass spectrometry experiments acquired over a mass range from 150 to 2200 m/z using an electrospray ionization source operated at a temperature of 200C and a spray voltage of 4.5 kV.

#### Peptide identification

Peptides were identified from the non-deuterated samples of p110γ alone or p110γ complexed with p101 or p84 using data-dependent acquisition following tandem MS/MS experiments (0.5 s precursor scan from 150-2000 m/z; twelve 0.25 s fragment scans from 150-2000 m/z). MS/MS datasets were analyzed using PEAKS7 (PEAKS), and peptide identification was carried out by using a false discovery based approach, with a threshold set to 1% using a database of of known contaminants found in Sf9 cells (Dobbs et al., 2020). The search parameters were set with a precursor tolerance of 20 ppm, fragment mass error 0.02 Da, charge states from 1-8, leading to a selection criterion of peptides that had a −10logP score of 21.7.

#### Mass Analysis of Peptide Centroids and Measurement of Deuterium Incorporation

HD-Examiner Software (Sierra Analytics) was used to automatically calculate the level of deuterium incorporation into each peptide. All peptides were manually inspected for correct charge state, correct retention time, appropriate selection of isotopic distribution, etc. Deuteration levels were calculated using the centroid of the experimental isotope clusters. Results are presented as relative levels of deuterium incorporation, with the only correction being applied correcting for the deuterium oxide percentage of the buffer utilized in the exchange (84.5% and 86.8%). Differences in exchange in a peptide were considered significant if they met all two of the following criteria: ≥5% change in exchange and ≥0.4 Da difference in exchange. The raw HDX data are shown in two different formats. To allow for visualization of differences across all peptides, we utilized number of deuteron difference (#D) plots (Fig 2B). These plots show the total difference in deuterium incorporation over the entire H/D exchange time course, with each point indicating a single peptide. Samples were only compared within a single experiment and were never compared to experiments completed at a different time with a different final D_2_O level. The data analysis statistics for all HDX-MS experiments are in Supplemental Table 2 according to the guidelines of (Masson et al., 2019). The mass spectrometry proteomics data have been deposited to the ProteomeXchange Consortium via the PRIDE partner repository (Perez-Riverol et al., 2019) with the dataset identifier PXD025207.

#### Lipid vesicle preparation

For kinase assays, lipid vesicles containing 5% brain phosphatidylinositol 4,5-bisphosphate (PIP2), 20% brain phosphatidylserine (PS), 35% egg-yolk phosphatidylethanolamine (PE), 10% egg-yolk phosphatidylcholine (PC), 25% cholesterol and 5% egg-yolk sphingomyelin (SM) were prepared by mixing the lipids dissolved in organic solvent. The solvent was evaporated in a stream of argon following which the lipid film was desiccated in a vacuum for 45 minutes. The lipids were resuspended in lipid buffer (20 mM HEPES pH 7.0, 100 mM NaCl and 10 % glycerol) and the solution was sonicated for 15 minutes. The vesicles were subjected to five freeze thaw cycles and extruded 11 times through a 100-nm filter (T&T Scientific: TT-002-0010). The extruded vesicles were sonicated again for 5 minutes, aliquoted and stored at −80°C.

#### In vitro lipid kinase assays

All lipid kinase activity assays employed the Transcreener ADP2 Fluorescence Intensity (FI) Assay (Bellbrook labs) which measures ADP production. For assays with nanobodies, PM-mimic vesicles (5% PIP2, 20% PS, 10% PC, 35% PE, 25% cholesterol, 5% SM) were used at a final concentration of 1 mg/mL, with ATP at a final concentration of 100 μM ATP and Gβγ/HRas at 1.5 μM final concentration were used. The protein solution at 2X final concentration was mixed with nanobody at 2X final concentration in a black 385 well microplate (Corning). 2 μL of the protein-nanobody solution was mixed with 2 μL substrate solution containing ATP, vesicles and Gβγ/HRas or Gβγ/HRas gel filtration buffer and the reaction was allowed to proceed for 60 minutes at 20°C. Final concentration of kinase was 1200-3500nM for all basal conditions with or without nanobody. For conditions with Gβγ (with or without nanobody), p110γ-p84: 400-1500 nM and p110γ-p101: 25-750 nM. For conditions with Ras (with or without nanobody), p110γ-p84: 1000-3000 nM and p110γ-p101: 750-3000 nM. For conditions with Gβγ and Ras (with or without nanobody), p110γ-p84: 120-3000 nM and p110γ-p101: 5-3000 nM. The final nanobody concentration was 6 μM. The reaction was stopped with 4 μL of 2X stop and detect solution containing Stop and Detect buffer, 8 nM ADP Alexa Fluor 594 Tracer and 93.7 μg/mL ADP2 Antibody IRDye QC-1 and incubated for 50 minutes. The fluorescence intensity was measured using a SpectraMax M5 plate reader at excitation 590 nm and emission 620 nm. This data was normalized was normalized against the measurements obtained for 100 μM ATP and 100 μM ADP. The % ATP turnover was interpolated from a standard curve (0.1-100 μM ADP). This was then used to calculate the specific activity of the enzyme.

#### Negative stain electron microscopy and image analysis

Purified p110γ-p101 in complex without (apo) or with nanobodies (NB1-PIK3R5, NB2-PIK3R5, NB5-PIK3CG) were adsorbed to glow discharged carbon coated grids at a concentration of 0.02 mg/mL for 30s and stained with uranyl formate. The stained specimens were examined using a Tecnai Spirit (apo, NB5-PIK3CG) or a Talos L120C (NB1-PIK3R5, NB2-PIK3R5) transmission electron microscope (ThermoFisher Scientific) operated at an accelerating voltage of 120 kV and equipped with an FEI Eagle 4K or Ceta charged-coupled-device (CCD) camera, respectively. For the apo-p110γ-p101 complex, 50 micrographs were acquired at a nominal magnification of 49,000x at a defocus of −1.2μm and binned by 2 to obtain a final pixel size of 4.67 Å/pixel. For the p110γ-p101-NB5-PIK3CG complex, 25 micrographs were acquired at a nominal magnification of 49,000x at a defocus of −1.2μm and binned twice to obtain a final pixel size of 4.67 Å/pixel. 80 micrographs of the p110γ-p101-NB2-PIK3R5 complex were acquired at a nominal magnification of 45,000x at a defocus of −1.2μm and binned twice to obtain a final pixel size of 4.53 Å/pixel. Finally, 50 micrographs of the p110γ-p101-NB1-PIK3R5 complex were acquired at a nominal magnification of 45,000x at a defocus of −1.2μm and binned twice to obtain a final pixel size of 4.53 Å/pixel. For each dataset, the contrast transfer function (CTF) of each micrograph was estimated using CTFFind4. 200 particles were manually picked then aligned to generate 2D class averages for template-based autopicking in Relion 3.0. These templates were then used to autopick 35629, 6425, 46848, and 23112 particles for the apo-, NB5-PIK3CG, NB3-PIK3R5, and NB1-PIK3R5-bound datasets, respectively, and extracted with a box size of 320Å. Particles were then subjected to 2D classification and the classes showing clear additional density for bound nanobodies when compared to the apo-complex were selected.

#### Cryo-EM Sample Preparation and Data Collection

C-Flat 2/2-T 300 mesh grids were glow discharged for 25s at 15mA using a Pelco easiGlow glow-discharger. 3μL of purified p110γ-p101 complex with or without bound nanobody was then applied to the grids at a concentration of 0.45 mg/ml. Grids were then prepared using a Vitrobot Mark IV (Thermo Fisher Scientific) by blotting for 1.5s at 4°C and 100% humidity with a blot force of −5 followed by plunge freezing in liquid ethane. Grids were screened for particle and ice quality at the UBC High Resolution Macromolecular Cryo-Electron Microscopy (HRMEM) facility using a 200kV Glacios TEM (Thermo Fisher Scientific) equipped with a Falcon 3EC DED. All datasets were then collected at the Pacific Northwest Cryo-EM Center (PNCC) using a Titan Krios equipped with a K3 DED and a BioQuantum K3 energy filter with a slit width of 20 eV (Gatan). For the apo p110γ-p101 complex, 6153 super-resolution movies were collected using SerialEM with a total dose of 50e^-^/Å^2^ over 50 frames at a physical pixel size of 1.079Å/pix, using a defocus range of −0.8 to −2μm. For the nanobody-bound p110γ-p101 complex, 6808 super-resolution movies were collected using SerialEM with a total dose of 36.4e^-^/Å^2^ over 50 frames at a physical pixel size of 1.059Å/pix, using a defocus range of −1 to −2.4μm.

#### Cryo-EM image analysis

All data processing was carried out using cryoSPARC v2.18+ unless otherwise specified. For the nanobody-bound p110γ-p101 complex dataset, patch motion correction using default settings was first applied to all movies to align the frames and Fourier crop the outputs by a factor of 2. The contrast transfer function (CTF) of the resulting micrographs was estimated using the patch CTF estimation job with default settings. 2D class averages from a previous dataset were low-pass filtered to 15Å and used as templates to auto-pick 3,762,631 particles, which were then extracted with a box size of 300 pixels. The particles were subjected to 2D classification with the 2D class re-center threshold set to 0.05, and a circular mask of 200Å. 2D class averages that were ice contamination or showed no features were discarded. The remaining 952,705 particles were next used for *ab initio* reconstruction and heterogenous refinement using 2 classes. 692,109 particles from the better 3D reconstruction were curated and any particles from micrographs with a CTF estimation worse than 3Å or total frame motion greater than 30Å were discarded. Per-particle local motion correction was then carried out on the remaining 662,855 particles. The particles were then used for *ab intio* reconstruction and heterogeneous refinement using 4 classes. 320,179 particles from the most complete class were used to carry out homogenous refinement using the 3D reconstruction for that class as a starting model, yielding a reconstruction with an overall resolution of 2.99Å based on the Fourier shell correlation (FSC) 0.143 criterion. The particles were further refined using local CTF refinement before being used for non-uniform refinement with simultaneous global CTF refinement, yielding a map with an overall resolution of 2.90Å. Finally, the map was subjected to a final non-uniform refinement using a mask enveloping the entire volume with the rotation fulcrum centered at the low-resolution nanobody-p101 interaction interface, producing the final map used for model building at a 2.89Å overall resolution.

For the apo p110γ-p101 complex dataset, full-frame motion correction using default settings was first applied to all movies to align the frames. The contrast transfer function (CTF) of the resulting micrographs was estimated using CTFFIND4 with default settings. 2D class averages from a previous dataset were low-pass filtered to 15Å and used as a template to auto-pick 4,792,176 particles, which were then down-sampled by 2 (resulting pixel size of 1.079 Å/pix) and extracted with a box size of 320 pixels.

The particles were subjected to multiple rounds of 2D classification with the 2D class re-center threshold set to 0.05, and a circular mask of 200Å. 2D class averages that did were ice contamination or did not align to high-resolution were then discarded. The remaining 1,285,510 particles were next subjected to patch CTF estimation and per-particle motion correction before being used for 2 more rounds of 2D classification. 731,169 particles which classified to “good” classes were then used for *ab initio* reconstruction and heterogenous refinement using 2 classes twice iteratively. 196,390 particles from the better 3D reconstruction were used to carry out homogenous refinement using the 3D reconstruction for that class as a starting model, yielding a reconstruction with an overall resolution of 3.49Å based on the Fourier shell correlation (FSC) 0.143 criterion. The map was further refined non-uniform refinement, yielding a map with an overall resolution of 3.36Å. The full details of the p110γ-p101 structure for this map is described in an a separate manuscript.

## Supplemental Figures and Tables

**Figure S1.**
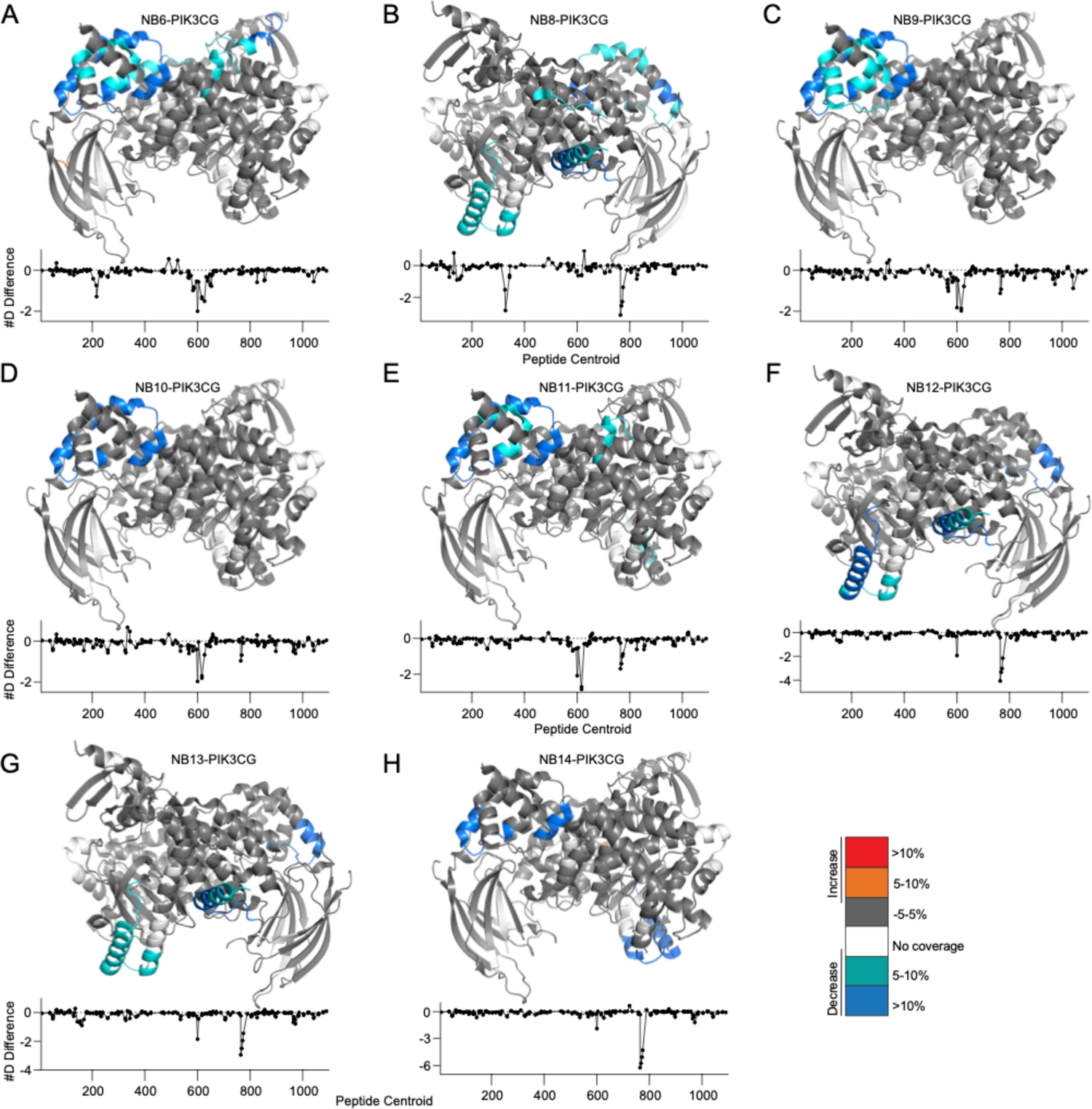
HDX-MS differences in p110γ on nanobody binding. **A.** HDX-MS differences in p110γ-p84 with the addition of NB6-PIK3CG mapped on a model of p110γ. The number of deuteron difference for all peptides analysed over the entire deuterium exchange time course is shown for p110γ. In panels **A-H**, peptides showing significant difference in deuterium exchange (>5%,>0.4 kDa) between conditions with and without nanobody are colored on the cartoon model. **B.** HDX-MS differences in p110γ-p84 with the addition of NB8-PIK3CG mapped on a model of p110γ. The number of deuteron difference for all peptides analyzed over the entire deuterium exchange time course is shown for p110γ. **C.** HDX-MS differences in p110γ-p84 with the addition of NB9-PIK3CG mapped on a model of p110γ. The number of deuteron difference for all peptides analyzed over the entire deuterium exchange time course is shown for p110γ. **D.** HDX-MS differences in p110γ-p84 with the addition of NB10-PIK3CG mapped on a model of p110γ. The number of deuteron difference for all peptides analyzed over the entire deuterium exchange time course is shown for p110γ. **E.** HDX-MS differences in p110γ-p84 with the addition of NB11-PIK3CG mapped on a model of p110γ. The number of deuteron difference for all peptides analyzed over the entire deuterium exchange time course is shown for p110γ. **F.** HDX-MS differences in p110γ-p84 with the addition of NB12-PIK3CG mapped on a model of p110γ. The number of deuteron difference for all peptides analyzed over the entire deuterium exchange time course is shown for p110γ. **G.** HDX-MS differences in p110γ-p84 with the addition of NB13-PIK3CG mapped on a model of p110γ. The number of deuteron difference for all peptides analyzed over the entire deuterium exchange time course is shown for p110γ. **H.** HDX-MS differences in p110γ-p84 with the addition of NB14-PIK3CG mapped on a model of p110γ. The number of deuteron difference for all peptides analyzed over the entire deuterium exchange time course is shown for p110γ.

**Figure S2.**
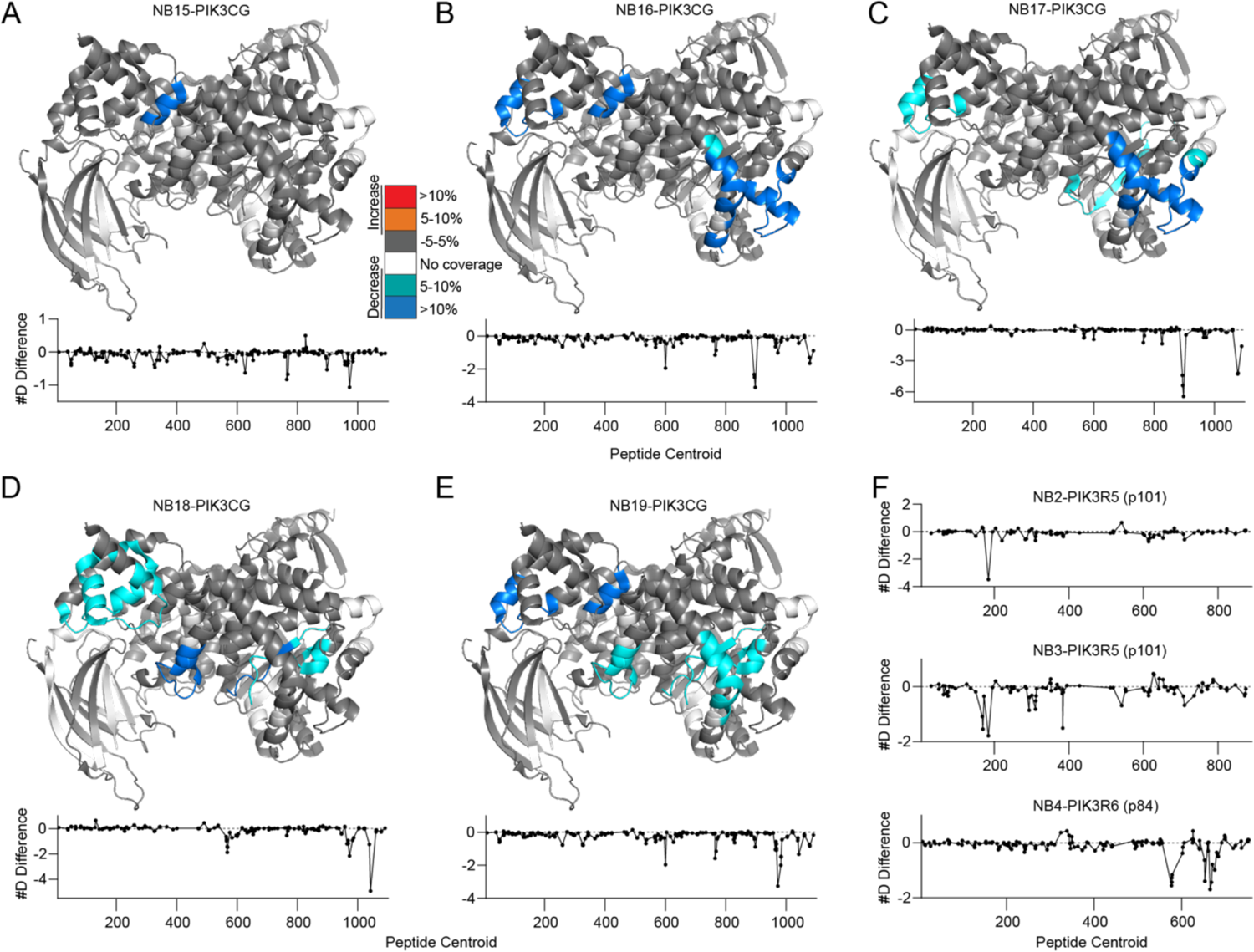
HDX-MS differences in p110γ, p101 and p84 on nanobody binding. **A.** HDX-MS differences in p110*γ*-p84 with the addition of NB15-PIK3CG mapped on a model of p110*γ*. The number of deuteron difference for all peptides analyzed over the entire deuterium exchange time course is shown for p110*γ*. For panels **A-E**, peptides showing significant difference in deuterium exchange (>5%,>0.4 kDa) between conditions with and without nanobody are colored on the cartoon model. **B.** HDX-MS differences in p110γ-p84 with the addition of NB16-PIK3CG mapped on a model of p110γ. The number of deuteron difference for all peptides analyzed over the entire deuterium exchange time course is shown for p110γ. **C.** HDX-MS differences in p110γ-p84 with the addition of NB17-PIK3CG mapped on a model of p110γ. The number of deuteron difference for all peptides analyzed over the entire deuterium exchange time course is shown for p110γ. **D.** HDX-MS differences in p110γ-p84 with the addition of NB18-PIK3CG mapped on a model of p110γ. The number of deuteron difference for all peptides analyzed over the entire deuterium exchange time course is shown for p110γ. **E.** HDX-MS differences in p110γ-p84 with the addition of NB19-PIK3CG mapped on a model of p110γ. The number of deuteron difference for all peptides analyzed over the entire deuterium exchange time course is shown for p110γ. **F.** The number of deuteron difference between conditions with and without nanobody for all peptides analyzed over the entire deuterium exchange time course is shown for p101 (NB2-PIK3R5 and NB3-PIK3R5) and p84 (NB4-PIK3R6).

**Table S1.** Nanobody protein specificity determined by pulldown assays, ΔTm induced by nanobody binding determined by DSF and peptides stabilized in HDX-MS.

| Internal Identifier | Nanobody | Protein Specificity | DSF stabilization ($\Delta T_m$ °C) | Regions with significant protection in HDX |
| --- | --- | --- | --- | --- |
| CA15452 | NB1-PIK3R5 | p101 | 0 | 176-193, 279-288, 605-623, 650-654 |
| CA15433 | NB2-PIK3R5 | p101 | 1 | 165-193, 289-298, 304-317, 378-389, 537-546 |
| CA15445 | NB3-PIK3R5 | p101 | 0 | 677-726, 816-830 |
| CA15424 | NB4-PIK3R6 | p84 | -0.5 | 650,655, 656-673 |
| CA15428 | NB5-PIK3CG | p110γ | 0.5 | 125-157, 623-630 |
| CA15423 | NB6-PIK3CG | p110γ | 3 | 196-221, 574-607, 611-630, 636-654, 849-861 |
| CA15468 | NB7-PIK3CG | p110γ | 3.3 | 196-211, 578-607, 611-630, 643-657, 816-838, 853-861, 1035-1050 |
| CA15460 | NB8-PIK3CG | p110γ | 0.3 | 138-164, 316-339, 593-607, 611-622, 748-782 |
| CA15441 | NB9-PIK3CG | p110γ | 0.3 | 557-578, 579-607, 611-635 |
| CA15442 | NB10-PIK3CG | p110γ | 0.2 | 593-607, 611-630 |
| CA15420 | NB11-PIK3CG | p110γ | 1.2 | 579-607, 611-630, 768-782, 849-861 |
| CA15403 | NB12-PIK3CG | p110γ | 1.1 | 138-164, 593-607, 748-782 |
| CA15416 | NB13-PIK3CG | p110γ | 0.6 | 138-164, 593-607, 748-782 |
| CA15454 | NB14-PIK3CG | p110γ | 0.1 | 593-607, 623-630, 748-782 |
| CA15430 | NB15-PIK3CG | p110γ | 0.4 | 623-630 |
| CA15457 | NB16-PIK3CG | p110γ | 0 | 593-607, 623-630, 888-907, 1072-1084, 1088-1092 |
| CA15470 | NB17-PIK3CG | p110γ | 0.2 | 593-607, 816-838, 888-910, 1072-1084, 1088-1092 |
| CA15481 | NB18-PIK3CG | p110γ | 1.5 | 557-601, 954-975, 961-992, 1035-1050 |
| CA15443 | NB19-PIK3CG | p110γ | 0.2 | 593-607, 623-630, 954-992, 1035-1050, 1072-1084 |

## References

1. Bannas, P., Hambach, J., Koch-Nolte, F., 2017. Nanobodies and Nanobody-Based Human Heavy Chain Antibodies As Antitumor Therapeutics. Front Immunol 8, 1603. doi:10.3389/fimmu.2017.01603

2. Baranova, E., Fronzes, R., Garcia-Pino, A., Van Gerven, N., Papapostolou, D., Péhau-Arnaudet, G., Pardon, E., Steyaert, J., Howorka, S., Remaut, H., 2012. SbsB structure and lattice reconstruction unveil Ca2+ triggered S-layer assembly. Nature 487, 119–122. doi:10.1038/nature11155

3. Beghein, E., Gettemans, J., 2017. Nanobody Technology: A Versatile Toolkit for Microscopic Imaging, Protein-Protein Interaction Analysis, and Protein Function Exploration. Front Immunol 8, 771. doi:10.3389/fimmu.2017.00771

4. Berkowitz, S.A., Engen, J.R., Mazzeo, J.R., Jones, G.B., 2012. Analytical tools for characterizing biopharmaceuticals and the implications for biosimilars. Nat Rev Drug Discov 11, 527–540. doi:10.1038/nrd3746

5. Bohnacker, T., Marone, R., Collmann, E., Calvez, R., Hirsch, E., Wymann, M., 2009. PI3Kgamma adaptor subunits define coupling to degranulation and cell motility by distinct PtdIns(3,4,5)P3 pools in mast cells. Sci Signal 2, ra27. doi:10.1126/scisignal.2000259

6. Buckles, T.C., Ohashi, Y., Tremel, S., McLaughlin, S.H., Pardon, E., Steyaert, J., Gordon, M.T., Williams, R.L., Falke, J.J., 2020. The G-Protein Rab5A Activates VPS34 Complex II, a Class III PI3K, by a Dual Regulatory Mechanism. Biophys. J. 119, 2205–2218. doi:10.1016/j.bpj.2020.10.028

7. Burke, J.E., Williams, R.L., 2015. Synergy in activating class I PI3Ks. Trends in Biochemical Sciences 40, 88–100. doi:10.1016/j.tibs.2014.12.003

8. Camps, M., Rückle, T., Ji, H., Ardissone, V., Rintelen, F., Shaw, J., Ferrandi, C., Chabert, C., Gillieron, C., Françon, B., Martin, T., Gretener, D., Perrin, D., Leroy, D., Vitte, P.-A., Hirsch, E., Wymann, M.P., Cirillo, R., Schwarz, M.K., Rommel, C., 2005. Blockade of PI3Kgamma suppresses joint inflammation and damage in mouse models of rheumatoid arthritis. Nat. Med. 11, 936–943. doi:10.1038/nm1284

9. De Groof, T.W.M., Bobkov, V., Heukers, R., Smit, M.J., 2019. Nanobodies: New avenues for imaging, stabilizing and modulating GPCRs. Mol Cell Endocrinol 484, 15–24. doi:10.1016/j.mce.2019.01.021

10. De Henau, O., Rausch, M., Winkler, D., Campesato, L.F., Liu, C., Cymerman, D.H., Budhu, S., Ghosh, A., Pink, M., Tchaicha, J., Douglas, M., Tibbitts, T., Sharma, S., Proctor, J., Kosmider, N., White, K., Stern, H., Soglia, J., Adams, J., Palombella, V.J., McGovern, K., Kutok, J.L., Wolchok, J.D., Merghoub, T., 2016. Overcoming resistance to checkpoint blockade therapy by targeting PI3Kγ in myeloid cells. Nature. doi:10.1038/nature20554

11. Deladeriere, A., Gambardella, L., Pan, D., Anderson, K.E., Hawkins, P.T., Stephens, L.R., 2015. The regulatory subunits of PI3Kγ control distinct neutrophil responses. Sci Signal 8, ra8. doi:10.1126/scisignal.2005564

12. Desmyter, A., Transue, T.R., Ghahroudi, M.A., Thi, M.H., Poortmans, F., Hamers, R., Muyldermans, S., Wyns, L., 1996. Crystal structure of a camel single-domain VH antibody fragment in complex with lysozyme. Nat. Struct. Biol. 3, 803–811. doi:10.1038/nsb0996-803

13. Dobbs, J.M., Jenkins, M.L., Burke, J.E., 2020. Escherichia coli and Sf9 Contaminant Databases to Increase Efficiency of Tandem Mass Spectrometry Peptide Identification in Structural Mass Spectrometry Experiments. J. Am. Soc. Mass Spectrom. 31, 2202–2209. doi:10.1021/jasms.0c00283

14. Domanska, K., Vanderhaegen, S., Srinivasan, V., Pardon, E., Dupeux, F., Marquez, J.A., Giorgetti, S., Stoppini, M., Wyns, L., Bellotti, V., Steyaert, J., 2011. Atomic structure of a nanobody-trapped domain-swapped dimer of an amyloidogenic beta2-microglobulin variant. Proc. Natl. Acad. Sci. U.S.A. 108, 1314–1319. doi:10.1073/pnas.1008560108

15. Fruman, D.A., Chiu, H., Hopkins, B.D., Bagrodia, S., Cantley, L.C., Abraham, R.T., 2017. The PI3K Pathway in Human Disease. Cell 170, 605–635. doi:10.1016/j.cell.2017.07.029

16. Gangadhara, G., Dahl, G., Bohnacker, T., Rae, R., Gunnarsson, J., Blaho, S., Öster, L., Lindmark, H., Karabelas, K., Pemberton, N., Tyrchan, C., Mogemark, M., Wymann, M.P., Williams, R.L., Perry, M.W.D., Papavoine, T., Petersen, J., 2019. A class of highly selective inhibitors bind to an active state of PI3Kγ. Nature Chemical Biology 15, 348–357. doi:10.1038/s41589-018-0215-0

17. García-Nafría, J., Lee, Y., Bai, X., Carpenter, B., Tate, C.G., 2018. Cryo-EM structure of the adenosine A2A receptor coupled to an engineered heterotrimeric G protein. Elife 7. doi:10.7554/eLife.35946

18. Gulati, S., Jin, H., Masuho, I., Orban, T., Cai, Y., Pardon, E., Martemyanov, K.A., Kiser, P.D., Stewart, P.L., Ford, C.P., Steyaert, J., Palczewski, K., 2018. Targeting G protein-coupled receptor signaling at the G protein level with a selective nanobody inhibitor. Nat Commun 9, 1996–15. doi:10.1038/s41467-018-04432-0

19. Hamers-Casterman, C., Atarhouch, T., Muyldermans, S., Robinson, G., Hamers, C., Songa, E.B., Bendahman, N., Hamers, R., 1993. Naturally occurring antibodies devoid of light chains. Nature 363, 446–448. doi:10.1038/363446a0

20. Hawkins, P.T., Stephens, L.R., 2015. PI3K signalling in inflammation. Biochim. Biophys. Acta 1851, 882–897. doi:10.1016/j.bbalip.2014.12.006

21. Huang, W., Manglik, A., Venkatakrishnan, A.J., Laeremans, T., Feinberg, E.N., Sanborn, A.L., Kato, H.E., Livingston, K.E., Thorsen, T.S., Kling, R.C., Granier, S., Gmeiner, P., Husbands, S.M., Traynor, J.R., Weis, W.I., Steyaert, J., Dror, R.O., Kobilka, B.K., 2015. Structural insights into µ-opioid receptor activation. Nature 524, 315–321. doi:10.1038/nature14886

22. Huo, J., Le Bas, A., Ruza, R.R., Duyvesteyn, H.M.E., Mikolajek, H., Malinauskas, T., Tan, T.K., Rijal, P., Dumoux, M., Ward, P.N., Ren, J., Zhou, D., Harrison, P.J., Weckener, M., Clare, D.K., Vogirala, V.K., Radecke, J., Moynié, L., Zhao, Y., Gilbert-Jaramillo, J., Knight, M.L., Tree, J.A., Buttigieg, K.R., Coombes, N., Elmore, M.J., Carroll, M.W., Carrique, L., Shah, P.N.M., James, W., Townsend, A.R., Stuart, D.I., Owens, R.J., Naismith, J.H., 2020. Neutralizing nanobodies bind SARS-CoV-2 spike RBD and block interaction with ACE2. Nature Structural & Molecular Biology 27, 846–854. doi:10.1038/s41594-020-0469-6

23. Irannejad, R., Tomshine, J.C., Tomshine, J.R., Chevalier, M., Mahoney, J.P., Steyaert, J., Rasmussen, S.G.F., Sunahara, R.K., El-Samad, H., Huang, B., Zastrow von, M., 2013. Conformational biosensors reveal GPCR signalling from endosomes. Nature 495, 534–538. doi:10.1038/nature12000

24. Kaneda, M.M., Messer, K.S., Ralainirina, N., Li, H., Leem, C.J., Gorjestani, S., Woo, G., Nguyen, A.V., Figueiredo, C.C., Foubert, P., Schmid, M.C., Pink, M., Winkler, D.G., Rausch, M., Palombella, V.J., Kutok, J., McGovern, K., Frazer, K.A., Wu, X., Karin, M., Sasik, R., Cohen, E.E.W., Varner, J.A., 2016. PI3Kγ is a molecular switch that controls immune suppression. Nature 539, 437–442. doi:10.1038/nature19834

25. Korotkov, K.V., Pardon, E., Steyaert, J., Hol, W.G.J., 2009. Crystal structure of the N-terminal domain of the secretin GspD from ETEC determined with the assistance of a nanobody. Structure/Folding and Design 17, 255–265. doi:10.1016/j.str.2008.11.011

26. Kozasa, T., Gilman, A.G., 1995. Purification of recombinant G proteins from Sf9 cells by hexahistidine tagging of associated subunits. Characterization of alpha 12 and inhibition of adenylyl cyclase by alpha z. J. Biol. Chem. 270, 1734–1741. doi:10.1074/jbc.270.4.1734

27. Kurig, B., Shymanets, A., Bohnacker, T., Prajwal, Brock, C., Ahmadian, M.R., Schaefer, M., Gohla, A., Harteneck, C., Wymann, M.P., Jeanclos, E., Nürnberg, B., 2009. Ras is an indispensable coregulator of the class IB phosphoinositide 3-kinase p87/p110gamma. Proc. Natl. Acad. Sci. U.S.A. 106, 20312–20317. doi:10.1073/pnas.0905506106

28. Laverty, D., Desai, R., Uchański, T., Masiulis, S., Stec, W.J., Malinauskas, T., Zivanov, J., Pardon, E., Steyaert, J., Miller, K.W., Aricescu, A.R., 2019. Cryo-EM structure of the human α1β3γ2 GABAA receptor in a lipid bilayer. Nature 565, 516–520. doi:10.1038/s41586-018-0833-4

29. Lawrence, M.S., Stojanov, P., Mermel, C.H., Robinson, J.T., Garraway, L.A., Golub, T.R., Meyerson, M., Gabriel, S.B., Lander, E.S., Getz, G., 2014. Discovery and saturation analysis of cancer genes across 21 tumour types. Nature 505, 495–501. doi:10.1038/nature12912

30. Luo, L., Wall, A.A., Tong, S.J., Hung, Y., Xiao, Z., Tarique, A.A., Sly, P.D., Fantino, E., Marzolo, M.-P., Stow, J.L., 2018. TLR Crosstalk Activates LRP1 to Recruit Rab8a and PI3Kγ for Suppression of Inflammatory Responses. Cell Rep 24, 3033–3044. doi:10.1016/j.celrep.2018.08.028

31. Luo, L., Wall, A.A., Yeo, J.C., Condon, N.D., Norwood, S.J., Schoenwaelder, S., Chen, K.W., Jackson, S., Jenkins, B.J., Hartland, E.L., Schroder, K., Collins, B.M., Sweet, M.J., Stow, J.L., 2014. Rab8a interacts directly with PI3Kγ to modulate TLR4-driven PI3K and mTOR signalling. Nat Commun 5, 4407. doi:10.1038/ncomms5407

32. Madsen, R.R., Vanhaesebroeck, B., 2020. Cracking the context-specific PI3K signaling code. Sci Signal 13, eaay2940. doi:10.1126/scisignal.aay2940

33. Maier, U., Babich, A., Nurnberg, B., 1999. Roles of non-catalytic subunits in gbetagamma-induced activation of class I phosphoinositide 3-kinase isoforms beta and gamma. J. Biol. Chem. 274, 29311–29317.

34. Manglik, A., Kobilka, B.K., Steyaert, J., 2017. Nanobodies to Study G Protein-Coupled Receptor Structure and Function. Annu. Rev. Pharmacol. Toxicol. 57, 19–37. doi:10.1146/annurev-pharmtox-010716-104710

35. Masson, G.R., Burke, J.E., Ahn, N.G., Anand, G.S., Borchers, C., Brier, S., Bou-Assaf, G.M., Engen, J.R., Englander, S.W., Faber, J., Garlish, R., Griffin, P.R., Gross, M.L., Guttman, M., Hamuro, Y., Heck, A.J.R., Houde, D., Iacob, R.E., Jørgensen, T.J.D., Kaltashov, I.A., Klinman, J.P., Konermann, L., Man, P., Mayne, L., Pascal, B.D., Reichmann, D., Skehel, M., Snijder, J., Strutzenberg, T.S., Underbakke, E.S., Wagner, C., Wales, T.E., Walters, B.T., Weis, D.D., Wilson, D.J., Wintrode, P.L., Zhang, Z., Zheng, J., Schriemer, D.C., Rand, K.D., 2019. Recommendations for performing, interpreting and reporting hydrogen deuterium exchange mass spectrometry (HDX-MS) experiments. Nat. Methods 16, 595–602. doi:10.1038/s41592-019-0459-y

36. Masson, G.R., Jenkins, M.L., Burke, J.E., 2017. An overview of hydrogen deuterium exchange mass spectrometry (HDX-MS) in drug discovery. Expert Opin Drug Discov 12, 981–994. doi:10.1080/17460441.2017.1363734

37. McMahon, C., Staus, D.P., Wingler, L.M., Wang, J., Skiba, M.A., Elgeti, M., Hubbell, W.L., Rockman, H.A., Kruse, A.C., Lefkowitz, R.J., 2020. Synthetic nanobodies as angiotensin receptor blockers. Proc. Natl. Acad. Sci. U.S.A. 117, 20284–20291. doi:10.1073/pnas.2009029117

38. McPhail, J.A., Burke, J.E., 2020. Drugging the Phosphoinositide 3-Kinase (PI3K) and Phosphatidylinositol 4-Kinase (PI4K) Family of Enzymes for Treatment of Cancer, Immune Disorders, and Viral/Parasitic Infections. Adv. Exp. Med. Biol. 1274, 203– 222. doi:10.1007/978-3-030-50621-6_9

39. Muyldermans, S., 2021. Applications of Nanobodies. Annu Rev Anim Biosci 9, 401–421. doi:10.1146/annurev-animal-021419-083831

40. Muyldermans, S., 2013. Nanobodies: natural single-domain antibodies. Annu. Rev. Biochem. 82, 775–797. doi:10.1146/annurev-biochem-063011-092449

41. Pacold, M.E., Suire, S., Perisic, O., Lara-Gonzalez, S., Davis, C.T., Walker, E.H., Hawkins, P.T., Stephens, L., Eccleston, J.F., Williams, R.L., 2000. Crystal structure and functional analysis of Ras binding to its effector phosphoinositide 3-kinase gamma. Cell 103, 931–943.

42. Pardon, E., Betti, C., Laeremans, T., Chevillard, F., Guillemyn, K., Kolb, P., Ballet, S., Steyaert, J., 2018. Nanobody-Enabled Reverse Pharmacology on G-Protein-Coupled Receptors. Angew. Chem. Int. Ed. Engl. 57, 5292–5295. doi:10.1002/anie.201712581

43. Pardon, E., Laeremans, T., Triest, S., Rasmussen, S.G.F., Wohlkonig, A., Ruf, A., Muyldermans, S., Hol, W.G.J., Kobilka, B.K., Steyaert, J., 2014. A general protocol for the generation of Nanobodies for structural biology. Nat Protoc 9, 674–693. doi:10.1038/nprot.2014.039

44. Patrucco, E., Notte, A., Barberis, L., Selvetella, G., Maffei, A., Brancaccio, M., Marengo, S., Russo, G., Azzolino, O., Rybalkin, S.D., Silengo, L., Altruda, F., Wetzker, R., Wymann, M.P., Lembo, G., Hirsch, E., 2004. PI3Kgamma modulates the cardiac response to chronic pressure overload by distinct kinase-dependent and - independent effects. Cell 118, 375–387. doi:10.1016/j.cell.2004.07.017

45. Perez-Riverol, Y., Csordas, A., Bai, J., Bernal-Llinares, M., Hewapathirana, S., Kundu, D.J., Inuganti, A., Griss, J., Mayer, G., Eisenacher, M., Pérez, E., Uszkoreit, J., Pfeuffer, J., Sachsenberg, T., Yilmaz, S., Tiwary, S., Cox, J., Audain, E., Walzer, M., Jarnuczak, A.F., Ternent, T., Brazma, A., Vizcaíno, J.A., 2019. The PRIDE database and related tools and resources in 2019: improving support for quantification data. Nucleic Acids Res. 47, D442–D450. doi:10.1093/nar/gky1106

46. Perino, A., Ghigo, A., Ferrero, E., Morello, F., Santulli, G., Baillie, G.S., Damilano, F., Dunlop, A.J., Pawson, C., Walser, R., Levi, R., Altruda, F., Silengo, L., Langeberg, L.K., Neubauer, G., Heymans, S., Lembo, G., Wymann, M.P., Wetzker, R., Houslay, M.D., Iaccarino, G., Scott, J.D., Hirsch, E., 2011. Integrating Cardiac PIP(3) and cAMP Signaling through a PKA Anchoring Function of p110gamma. Mol. Cell 42, 84–95. doi:10.1016/j.molcel.2011.01.030

47. Rasmussen, S.G.F., Choi, H.-J., Fung, J.J., Pardon, E., Casarosa, P., Chae, P.S., DeVree, B.T., Rosenbaum, D.M., Thian, F.S., Kobilka, T.S., Schnapp, A., Konetzki, I., Sunahara, R.K., Gellman, S.H., Pautsch, A., Steyaert, J., Weis, W.I., Kobilka, B.K., 2011a. Structure of a nanobody-stabilized active state of the β(2) adrenoceptor. Nature 469, 175–180. doi:10.1038/nature09648

48. Rasmussen, S.G.F., DeVree, B.T., Zou, Y., Kruse, A.C., Chung, K.Y., Kobilka, T.S., Thian, F.S., Chae, P.S., Pardon, E., Calinski, D., Mathiesen, J.M., Shah, S.T.A., Lyons, J.A., Caffrey, M., Gellman, S.H., Steyaert, J., Skiniotis, G., Weis, W.I., Sunahara, R.K., Kobilka, B.K., 2011b. Crystal structure of the β2 adrenergic receptor-Gs protein complex. Nature 477, 549–555. doi:10.1038/nature10361

49. Rathinaswamy, M.K., Gaieb, Z., Fleming, K.D., Borsari, C., Harris, N.J., Moeller, B.J., Wymann, M.P., Amaro, R.E., Burke, J.E., 2021. Disease related mutations in PI3Kγ disrupt regulatory C-terminal dynamics and reveal a path to selective inhibitors. Elife 10. doi:10.7554/eLife.64691

50. Rostislavleva, K., Soler, N., Ohashi, Y., Zhang, L., Pardon, E., Burke, J.E., Masson, G.R., Johnson, C., Steyaert, J., Ktistakis, N.T., Williams, R.L., 2015. Structure and flexibility of the endosomal Vps34 complex reveals the basis of its function on membranes. Science 350, aac7365. doi:10.1126/science.aac7365

51. Ruprecht, J.J., King, M.S., Zögg, T., Aleksandrova, A.A., Pardon, E., Crichton, P.G., Steyaert, J., Kunji, E.R.S., 2019. The Molecular Mechanism of Transport by the Mitochondrial ADP/ATP Carrier. Cell 176, 435–447.e15. doi:10.1016/j.cell.2018.11.025

52. Rynkiewicz, N.K., Anderson, K.E., Suire, S., Collins, D.M., Karanasios, E., Vadas, O., Williams, R., Oxley, D., Clark, J., Stephens, L.R., Hawkins, P.T., 2020. Gβγ is a direct regulator of endogenous p101/p110γ and p84/p110γ PI3Kγ complexes in mouse neutrophils. Sci Signal 13. doi:10.1126/scisignal.aaz4003

53. Scholler, P., Nevoltris, D., de Bundel, D., Bossi, S., Moreno-Delgado, D., Rovira, X., Møller, T.C., Moustaine, El, D., Mathieu, M., Blanc, E., McLean, H., Dupuis, E., Mathis, G., Trinquet, E., Daniel, H., Valjent, E., Baty, D., Chames, P., Rondard, P., Pin, J.-P., 2017. Allosteric nanobodies uncover a role of hippocampal mGlu2 receptor homodimers in contextual fear consolidation. Nat Commun 8, 1967–12. doi:10.1038/s41467-017-01489-1

54. Schubert, A.F., Gladkova, C., Pardon, E., Wagstaff, J.L., Freund, S.M.V., Steyaert, J., Maslen, S.L., Komander, D., 2017. Structure of PINK1 in complex with its substrate ubiquitin. Nature 552, 51–56. doi:10.1038/nature24645

55. Shymanets, A., Prajwal, P., Bucher, K., Beer-Hammer, S., Harteneck, C., Nürnberg, B., 2013. p87 and p101 subunits are distinct regulators determining class IB PI3K specificity. J. Biol. Chem. doi:10.1074/jbc.M113.508234

56. Shymanets, A., Prajwal, Vadas, O., Czupalla, C., LoPiccolo, J., Brenowitz, M., Ghigo, A., Hirsch, E., Krause, E., Wetzker, R., Williams, R.L., Harteneck, C., Nürnberg, B., 2015. Different inhibition of Gβγ-stimulated class IB phosphoinositide 3-kinase (PI3K) variants by a monoclonal antibody. Specific function of p101 as a Gβγ-dependent regulator of PI3Kγ enzymatic activity. Biochem. J. 469, 59–69. doi:10.1042/BJ20150099

57. Smirnova, I., Kasho, V., Jiang, X., Pardon, E., Steyaert, J., Kaback, H.R., 2015. Transient conformers of LacY are trapped by nanobodies. Proc. Natl. Acad. Sci. U.S.A. 112, 13839–13844. doi:10.1073/pnas.1519485112

58. Staus, D.P., Strachan, R.T., Manglik, A., Pani, B., Kahsai, A.W., Kim, T.H., Wingler, L.M., Ahn, S., Chatterjee, A., Masoudi, A., Kruse, A.C., Pardon, E., Steyaert, J., Weis, W.I., Prosser, R.S., Kobilka, B.K., Costa, T., Lefkowitz, R.J., 2016. Allosteric nanobodies reveal the dynamic range and diverse mechanisms of G-protein-coupled receptor activation. Nature 535, 448–452. doi:10.1038/nature18636

59. Steeland, S., Vandenbroucke, R.E., Libert, C., 2016. Nanobodies as therapeutics: big opportunities for small antibodies. Drug Discovery Today 21, 1076–1113. doi:10.1016/j.drudis.2016.04.003

60. Stephens, L.R., Eguinoa, A., Erdjument-Bromage, H., Lui, M., Cooke, F., Coadwell, J., Smrcka, A.S., Thelen, M., Cadwallader, K., Tempst, P., Hawkins, P.T., 1997. The Gbg sensitivity of a PI3K is dependent upon a tightly associated adaptor, p101. Cell 89, 105–114.

61. Uchański, T., Pardon, E., Steyaert, J., 2020. Nanobodies to study protein conformational states. Curr. Opin. Struct. Biol. 60, 117–123. doi:10.1016/j.sbi.2020.01.003

62. Vadas, O., Burke, J.E., Zhang, X., Berndt, A., Williams, R.L., 2011. Structural basis for activation and inhibition of class I phosphoinositide 3-kinases. Sci Signal 4, 1–13. doi:10.1126/scisignal.2002165

63. Vadas, O., Dbouk, H.A., Shymanets, A., Perisic, O., Burke, J.E., Abi Saab, W.F., Khalil, B.D., Harteneck, C., Bresnick, A.R., Nürnberg, B., Backer, J.M., Williams, R.L., 2013. Molecular determinants of PI3Kγ-mediated activation downstream of G-protein-coupled receptors (GPCRs). Proc. Natl. Acad. Sci. U.S.A. 110, 18862– 18867. doi:10.1073/pnas.1304801110

64. Westfield, G.H., Rasmussen, S.G.F., Su, M., Dutta, S., DeVree, B.T., Chung, K.Y., Calinski, D., Vélez-Ruiz, G., Oleskie, A.N., Pardon, E., Chae, P.S., Liu, T., Li, S., Woods, V.L., Steyaert, J., Kobilka, B.K., Sunahara, R.K., Skiniotis, G., 2011. Structural flexibility of the G alpha s alpha-helical domain in the beta2-adrenoceptor Gs complex. Proc. Natl. Acad. Sci. U.S.A. 108, 16086–16091. doi:10.1073/pnas.1113645108

65. Wingler, L.M., McMahon, C., Staus, D.P., Lefkowitz, R.J., Kruse, A.C., 2019. Distinctive Activation Mechanism for Angiotensin Receptor Revealed by a Synthetic Nanobody. Cell 176, 479–490.e12. doi:10.1016/j.cell.2018.12.006

66. Wrapp, D., De Vlieger, D., Corbett, K.S., Torres, G.M., Wang, N., Van Breedam, W., Roose, K., van Schie, L., VIB-CMB COVID-19 Response Team, Hoffmann, M., Pöhlmann, S., Graham, B.S., Callewaert, N., Schepens, B., Saelens, X., McLellan, J.S., 2020. Structural Basis for Potent Neutralization of Betacoronaviruses by Single-Domain Camelid Antibodies. Cell 181, 1004–1015.e15. doi:10.1016/j.cell.2020.04.031

